# The topography of gene tree topology space in a plant genus with a legacy of recent polyploidy and introgression

**DOI:** 10.1101/2024.09.27.615508

**Authors:** Jacob B. Landis, Andrew D. Farmer, Lucio Garcia, Racella McNair, Mariana Franco Ruiz, Qingli Liu, Jeff J. Doyle

**Affiliations:** School of Integrative Plant Science, Section of Plant Biology and the L.H. Bailey Hortorium, Cornell University, Ithaca NY 14853, USA; National Center for Genome Resources, Santa Fe NM 87505, USA; Seeds Research, Syngenta Crop Protection, LLC, Research Triangle Park, Durham NC 27709, USA

**Keywords:** clustering, *Glycine*, incongruence, maximum likelihood, species tree, pseudo-orthology, polyploidy

## Abstract

The eukaryotic genome has been described as a collection of different histories; for any set of taxa one of these histories is the record of cladogenic events that together comprise the species tree. Among the other histories expected to occur are those attributable to deep coalescence/lineage sorting; to biological causes such as introgression and horizontal transfer; or to pseudo-orthology, long branch attraction, and other technical issues. Gene tree topology space is the portion of tree space occupied by the gene trees reconstructed for a particular dataset of sampled genetic loci. Because coalescent theory predicts that the species tree topology will generally be the most frequent among gene trees, a reasonable expectation is that there will be a peak in gene tree topology space at the species tree topology, with secondary peaks present due to trees tracking other histories. Gene tree topology space in the small (∼30 species, including the cultivated soybean) legume genus, *Glycine* should not only have signals from the species tree and from lineage sorting, but also from a likely introgression event that created incongruence between the plastid and nuclear genomes. Additionally, *Glycine* is the product of a relatively recent (<13 million years) whole genome duplication, raising the possibility of pseudo-orthology. We explored this space using a set of 2389 nuclear genes and representative accessions from a 570-taxon concatenation tree, reconstructing gene trees for all nuclear loci and from complete plastid genomes and partial mitochondrial genomes. Species trees (ASTRAL) and maximum likelihood (ML) concatenation trees were congruent for a 61-taxon dataset but were incongruent with organellar genome trees. Gene tree topology space was flat: No topology was represented by more than one gene tree. This was also true for a reduced dataset of 27 taxa; only when the dataset was reduced to six ingroup taxa were multiple gene trees having the species tree topology observed, along with a topology congruent with the chloroplast genome topology, presumably representing nuclear loci introgressed along with the plastome. Clustering failed to identify any regional differentiation of gene tree topology space populated by loci with similar topologies. Pseudo-orthology did not contribute meaningfully to incongruence, in agreement with recent modeling work that minimizes concerns about this phenomenon. Clearly, different genes have different historical signals, but these signals are complex and exist at the level of clades within trees rather than as entire gene trees.

## Introduction

“Tree space” is a familiar concept in phylogenetics. Defined most simply as the set of possible tree topologies for a given number of operational taxonomic units (OTUs, taxa), it is searched to identify optimal solutions for particular datasets, and the resulting distribution of optimality scores is commonly conceptualized as being structured, with “islands” of trees possessing equal scores separated by suboptimal regions (Maddison 1991). The current paradigm of systematics explicitly distinguishes gene trees from species trees (Nei 1987), and uses information from multiple genes to estimate the underlying species tree (Edwards 2009; Bravo et al. 2019). Thus, for any phylogenomic analysis, there is a “species tree topology space” and a “gene tree topology space”. Phylogenomic studies identify a single tree; thus “species tree topology space” is the size of tree space for the number of OTUs sampled, populated at one topology by a single tree.

The corresponding “gene tree topology space” is far more complex. The species tree topology is reconstructed from alleles sampled from individual organisms; in sexually reproducing species these organisms “can be viewed as collections of genomic regions with different histories” (Rosenberg and Nordborg 2002). These “genomic regions” are “genes” in the coalescent sense (c-genes; Doyle 1995) defined as the minimal historical units freely recombining with one another but internally non-recombining (e.g., Springer and Gatesy 2018; Kubatko 2019; Doyle 2022) that may be larger or smaller than genes in the functional, molecular sense. Lack of internal recombination allows their histories to be depicted as acyclic graphs (i.e., gene trees). This recombinational freedom from one another allows genes to be bearers of “different histories”, whose often incongruent trees populate gene tree topology space. In a phylogenomic analysis that produces fully resolved gene trees for each of a set of 100 genetically unlinked loci sampled from 10 individuals, there can be at most 100 topologies represented by a gene tree among the ∼34 million possible topologies of tree space (Felsenstein 1978); if each locus had a different topology this would result in a flat landscape in gene tree topology space. Congruence among loci would result in topologies being represented by more than one locus, producing peaks in the gene tree topology landscape.

Population genetic theory predicts that for most speciation events the majority of gene tree histories will track the history of the species to which the sampled individuals belong, but that a minority of loci will have gene tree topologies incongruent with the species tree due to deep coalescence. This null hypothesis of incomplete lineage sorting (ILS) should result in gene tree topology space being structured, with a dominant species tree peak and smaller satellite peaks at alternative topologies, as is commonly illustrated for the simplest case of four taxa (e.g., Nei 1987; Maddison 1997). The sizes of ILS peaks relative to the species tree peak are determined by effective population sizes and times between speciation events.

Other “different histories” alter this neutral expectation of the gene tree topology landscape. Hybridization and introgression are pervasive across the tree of life (Mallet et al. 2016; Goulet et al. 2017), and horizontal transfer occurs not only in bacteria but in many eukaryotes (Van Etten and Bhattacharya 2020) including some angiosperm lineages (e.g., Poales: Dunning et al. 2019; Raimondeau et al. 2023). These processes do not produce a conflicting species tree, but instead transform the species tree’s acyclic graph into a cyclic graph (network). In this process, gene tree OTUs (alleles) are re-labeled with different species membership; no novel gene tree topologies are created, but transferred loci join one of the existing bins of loci whose topologies conflict with the species tree due to ILS. This asymmetric addition to the otherwise symmetric distribution of ILS topologies is the basis for tests of introgression (Durand et al. 2011).

There are many other sources of incongruence among gene trees (Wendel and Doyle 1998; Steenwyk et al. 2023) that can produce spurious signals in the gene tree topology landscape. Because orthologous genes trace their divergence to speciation events, whereas paralogues trace their origins to duplications (Fitch 1970), there is longstanding concern that conflation of paralogues and orthologues will lead to gene trees that do not reflect species relationships (Goodman et al. 1979; Doyle 1992; Maddison 1997; Edwards 2009; Struck 2013; Yang and Smith 2014). Of greatest concern is the potential for “pseudo-orthology” (Koonin 2005) or “artifactual orthologues” (Frost et al. 2024), in which the most recent common ancestor (MRCA) of the ingroup experienced a gene duplication that was followed by reciprocal loss of one paralogue in all ingroup species, with a different paralogue being lost by some species. A good gene tree is produced which is positively misleading as a species tree. Pseudo-orthology is of particular concern for plants, where polyploidy is prevalent, involving cycles of whole genome duplication that alternate with massive gene loss (fractionation) during diploidization, and generate complex gene families (Wendel 2015; Panchy et al. 2016; One Thousand Plant Transcriptomes Initiative 2019). Although pseudo-orthology may not be a major problem for most phylogenomic datasets, the impact of polyploidy in generating pseudo-orthologues has not been explored extensively (Smith and Hahn 2021, 2022; Smith et al. 2022) and remains a concern for some (Frost et al. 2024).

Overall, the gene tree topology space landscape for plants could be visualized as discrete peaks rising above the noise of random error in gene tree reconstruction. The major peak would be the species tree topology, with secondary peaks populated by topologies generated by deeply coalescing loci, introgressed loci, and loci where sampling a mixture of paralogues and orthologues has led to pseudo-orthology. But is this really what gene tree topology space looks like? The leguminous flowering plant genus, *Glycine*, is an excellent group for exploring this question. *Glycine* comprises around 30 species, including the cultivated soybean (*G. max*) and its wild progenitor (*G. soja*), annual species that together constitute subgenus *Soja*, native to eastern Asia. The remaining species are perennials belonging to subgenus *Glycine* and are mostly native to Australia (Sherman-Broyles et al. 2014b). Most *Glycine* species are functionally diploid at 2*n* = 40 (others are more recent tetraploids, 2*n* = 78, 80), but this high chromosome number relative to their mostly 2*n* = 20 or 22 generic relatives suggested ancestral polyploidy, as did the soybean genetic map and distribution of duplicate gene divergence times (Schlueter et al. 2004; Shoemaker et al. 2006). Because of its economic importance, soybean was an early target of whole genome sequencing (Schmutz et al. 2010), which showed that although the current haploid set of 20 chromosomes originated as two sets of 10 chromosomes with similar or identical synteny relationships, each chromosome is now a mosaic comprising blocks of homoeologous genes estimated to have diverged around 13 million years ago (MYA; Egan and Doyle 2010; Schmutz et al. 2010). This likely sets a maximum age for the polyploidy event itself (Doyle and Egan 2010), following which, in addition to extensive chromosomal rearrangements, around 25% of soybean WGD gene pairs lost one homoeologous sequence (Schmutz et al. 2010). Complete genomes have been sequenced for six diploid perennial species and one tetraploid, and show a high level of conservation in gene content and synteny with soybean (Chang et al. 2013; Zhuang et al. 2022). The genus thus represents a good study system for assessing the impact of pseudo-orthologues on phylogeny reconstruction.

As a potential resource for soybean improvement, subgenus *Glycine* has been extensively collected, facilitating decades of biosystematic studies, including DNA-based phylogenies beginning in the late 1980’s (reviewed in Sherman-Broyles et al. 2014b) and continuing into the phylogenomics era (e.g., Bombarely et al. 2014; Sherman-Broyles et al. 2014a, 2017). The morphologically and ecologically distinctive species, *G. falcata*, is supported as sister to the remaining perennial species in nuclear phylogenies (Doyle et al. 1999; Sherman-Broyles et al. 2014a), but molecular phylogenetic studies of the chloroplast genome (plastome) have suggested a very different placement for the species (Doyle et al. 1990a, 1990b; Sherman-Broyles et al. 2014b). Moreover, such cytonuclear incongruence was found to be the rule rather than the exception among reproductively compatible perennial *Glycine* species (“genome groups” of Hymowitz et al. 1998), and additional discordance was also found among nuclear genes, likely due to both deep coalescence and introgression (Doyle et al. 2004).

We generated genome resequencing data for 570 accessions of subgenus *Glycine* to identify single nucleotide polymorphisms (SNPs) from the nuclear and mitochondrial genomes, and assembled complete chloroplast genomes (plastomes). We sample from this large dataset to obtain a species tree for *Glycine* and phylogenetic hypotheses for their cytoplasmic genomes. We explore gene tree topology space for several *Glycine* datasets, finding that it is not dominated by any locus-specific phylogenetic signals produced either by the species tree or by competing signals from ILS or introgression. Leveraging synteny information from fully sequenced *Glycine* genomes, we also show that recent polyploidy and diploidization has not led to an accumulation of pseudo-orthologs.

## Methods and Materials

### Taxonomic sampling

Based on previous studies reporting crossing relationships and variation at individual nuclear and plastid loci (summarized above), and preliminary analyses of >500 accessions for the nuclear gene and whole plastome sequences described below (Supplemental Figure 1), a set of 61 accessions (Dataset 1: 61 perennials + *G. max* + *Amphicarpaea edgeworthii*) representing the diversity of the subgenus was constructed (Table 1). Two successively smaller subsets of these accessions (Dataset 2 - 27 accessions; Dataset 3 - six accessions from Zhuang et al. 2022) were used for exploration of gene tree topology space. Three datasets were designed to investigate the degree to which the utility of a gene in the taxonomically broader study (Dataset 2) predicted utility in more taxonomically focused datasets; each of these datasets consisted of the same number of OTUs as Dataset 2, and focused on: the A-genome (Dataset 4), the B-genome (Dataset 5), and the *G. tomentella* s.l. complex (D-, E-, H-, Ha-, and I-genomes; Dataset 6).

**Table 1.**
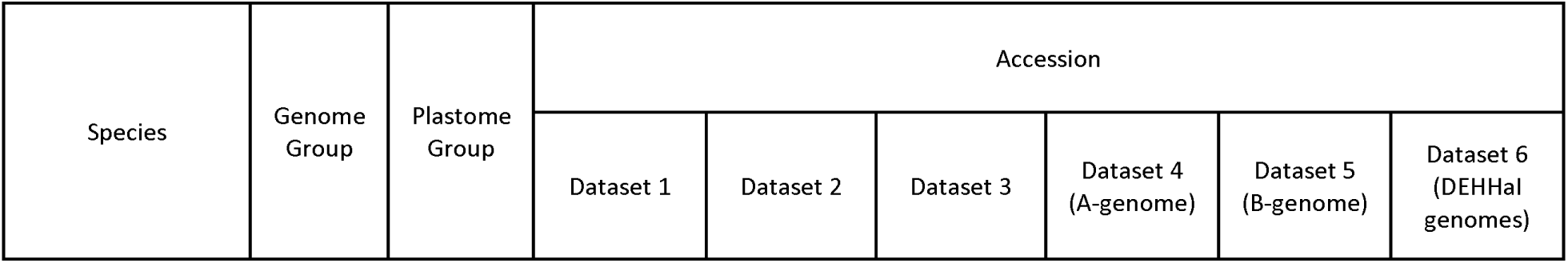
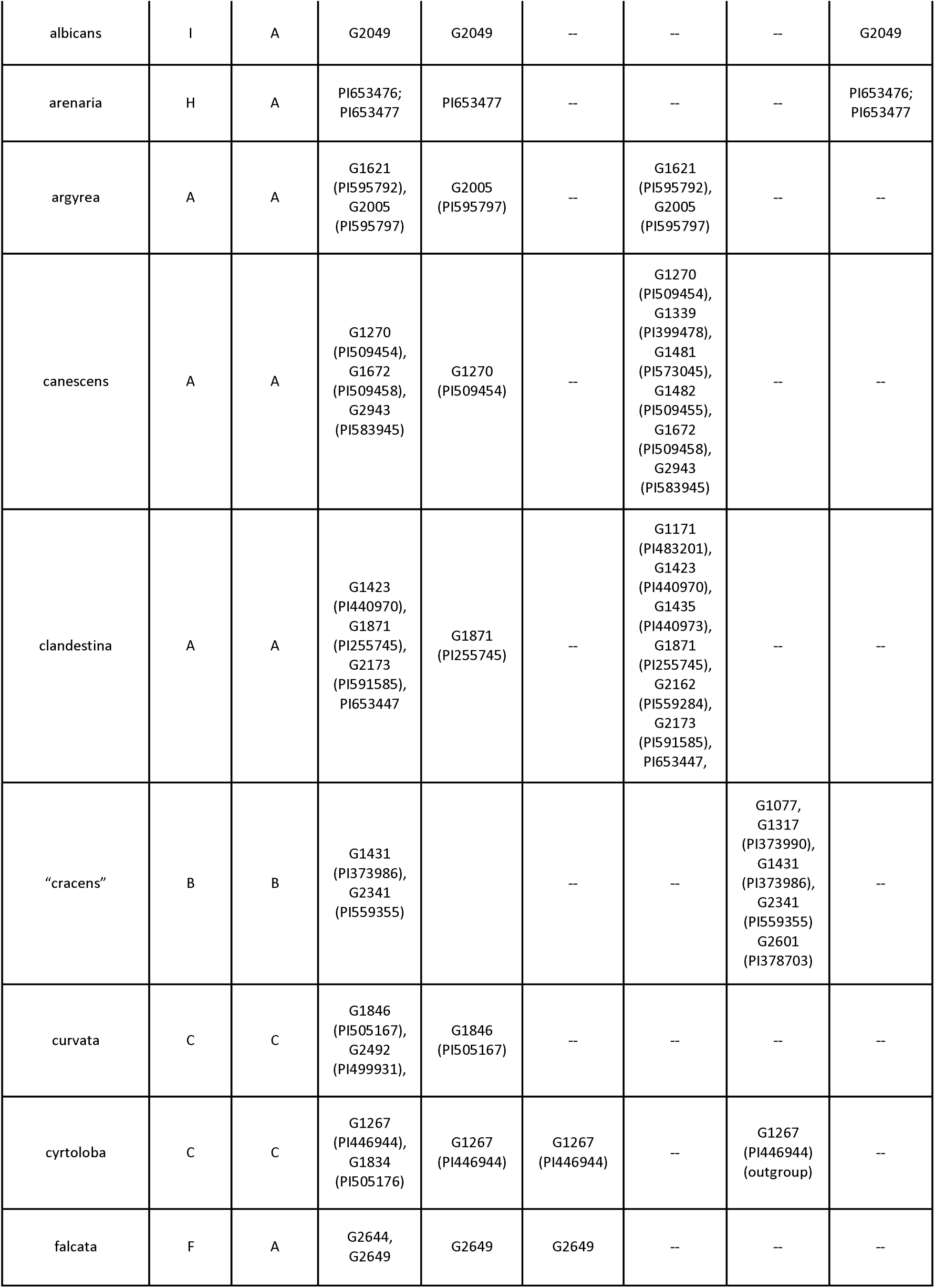
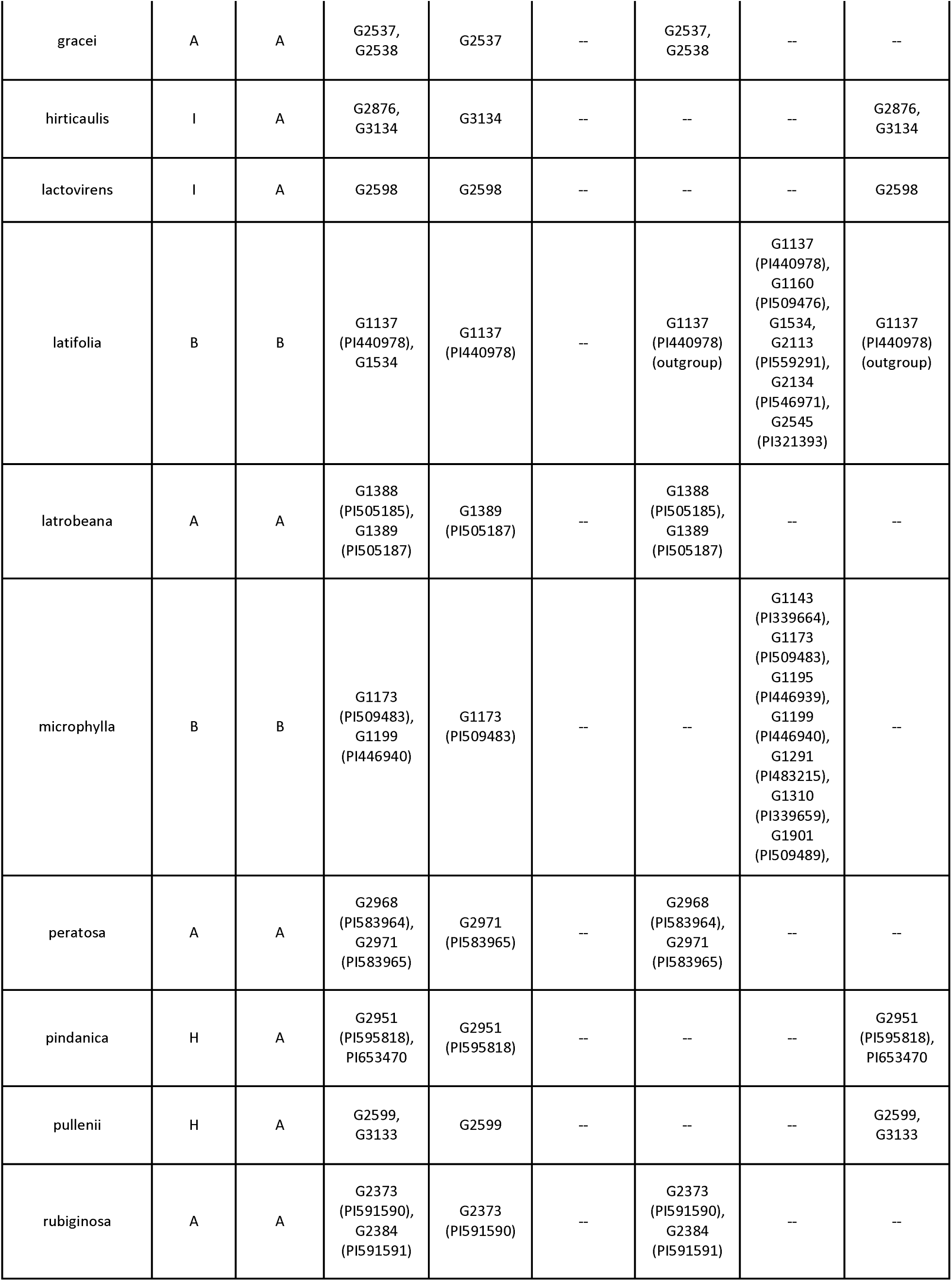
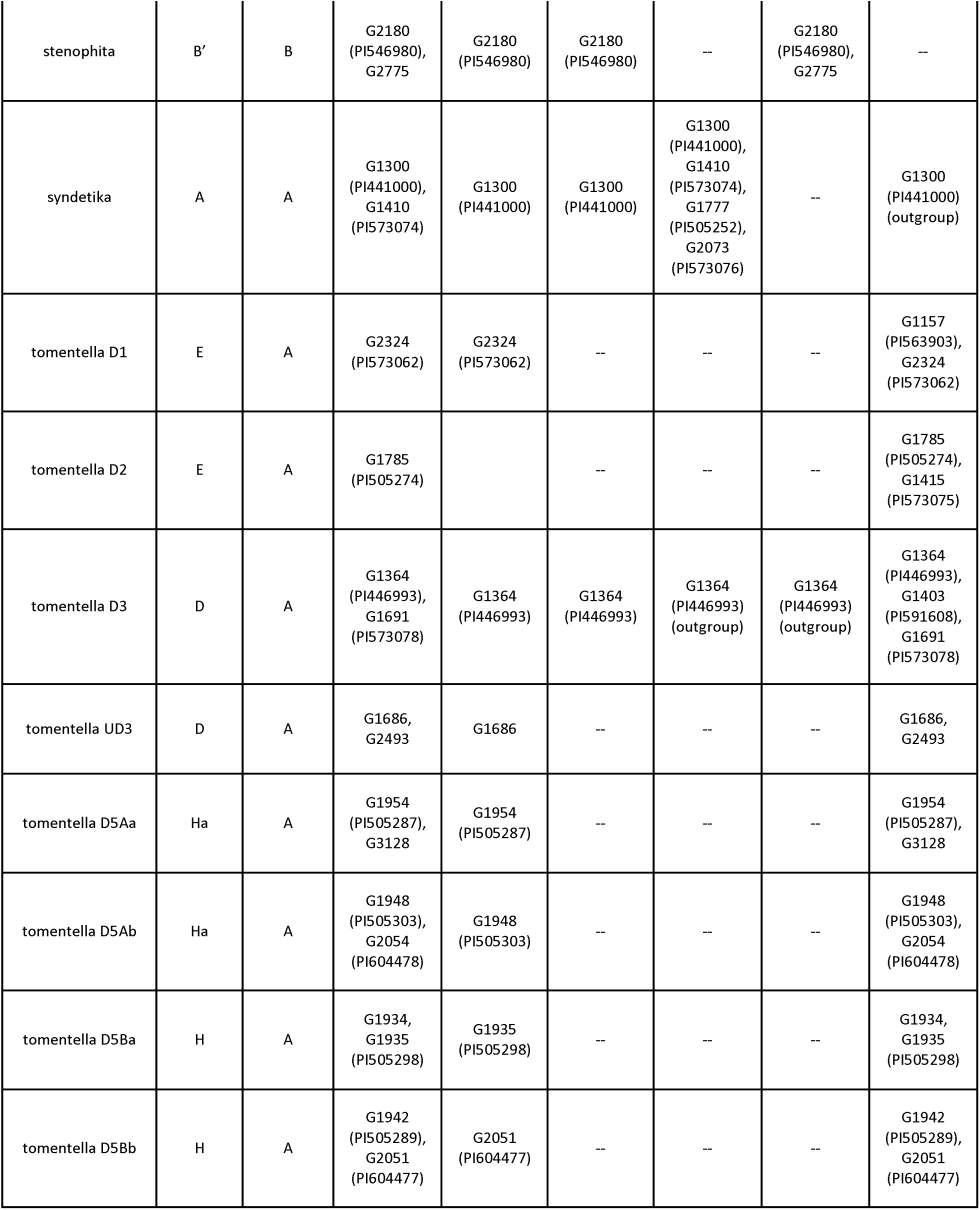
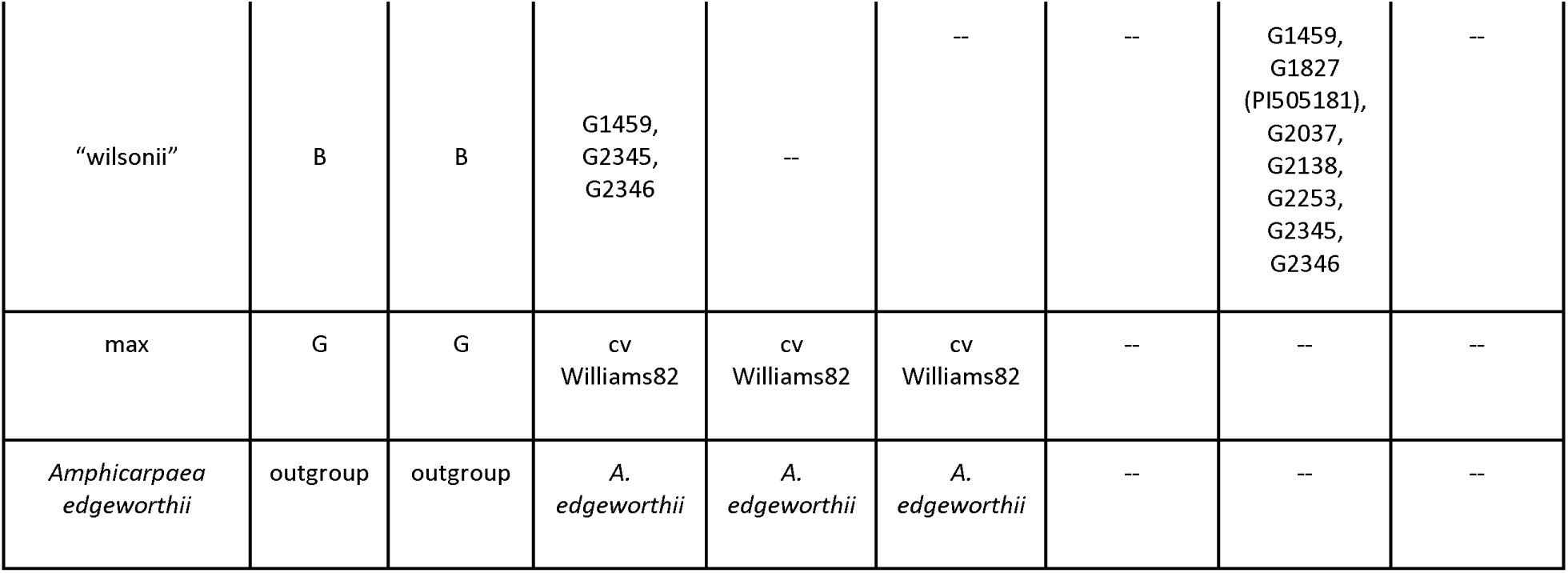
Accessions used for the different datasets including the unique identifier from the CSIRO (G-number) or USDA (PI-number).

### Whole-genome short-read sequencing

DNA was extracted from accessions using seed stocks from the CSIRO (Commonwealth Scientific and Industrial Research Organization) or USDA collections (Table 1) with CTAB (Doyle and Doyle 1987), followed by normalization to 15ng/µl. Sequencing libraries were prepared with 30 ng of normalized DNA using the NEBNext® Ultra™ II FS DNA Library Prep Kit for Illumina (New England BioLabs, Ipswich, Massachusetts, USA). Custom 8 bp barcodes were added for pooling, with up to ten samples in each pool, followed by cleaning with magnetic beads included in the library preparation kit. Each pool was run on one lane of Illumina HiSeqX (Illumina, San Diego, California, USA) with a 2×150 run yielding 150 gigabases. This resulted in a range from 8-21 gigabases for each sample.

### Nuclear gene sequences

The predicted proteins from an inhouse reference nuclear genome of *G. canescens* (PI599905) with the addition of *G. max* organellar genomes added from NCBI RefSeq (NC_007942 for the chloroplast genome and NC_020455 for the mitochondrial genome) were run through BUSCO v5.2.2 (Manni et al. 2021) using the embryophyta odb10 library resulting in 97.6% complete BUSCOS, with 37.6% being single copy and 60.0% duplicated, 1.4% fragmented, and 1.0% missing for the 1614 available BUSCO genes, corresponding to 2565 distinct loci in the reference genome. Of these, 2388 were retained for phylogenetic analyses plus the Histone H3D locus that has been previously used for phylogenetic inference in *Glycine* (Doyle et al. 1996; Pfeil et al. 2006; Landis and Doyle 2023). A fasta reference file for retained loci is available on Dryad (identifier provided upon acceptance). Paired Illumina reads were mapped to the reference loci using the preset short read parameter (-x sr) in minimap2 v2.17-r974-dirty (Li 2018). Genome-wide variant calls were made using freebayes v1.0.2-16-gd466dde (Garrison and Marth 2012) with the following parameters: --min-mapping-quality 4 --use-best-n-alleles 6 --no-population-priors --no-complex.

### Complete plastome sequences

Chloroplast genomes were assembled using getOrganelle v1.7.7.0 (Jin et al. 2020) using default parameters. The assembled plastomes were annotated using the web version of Chloë (https://chloe.plantenergy.edu.au/) with the first inverted repeat removed manually in AliView v1.28 (Larsson 2014). Pruned assemblies were aligned using progressiveMauve v.snapshop_2015_02_13 (Darling et al. 2010). The resulting alignment was manually inspected in AliView and local blocks were realigned using MAFFT v7.490 (Katoh and Standley 2013). A maximum likelihood tree was inferred using IQtree2 v.2.2.7 (Minh et al. 2020) with the GTR+G+F0 model of molecular evolution and 1000 bootstrap replicates.

### Mitochondrial genome sequences

SNPs were called against the *G. max* reference mitochondrial sequence (NC_020455) using FreeBayes v1.0.2-16-gd466dde (Garrison and Marth 2012). The reference sequence was included as an additional accession in the VCF file using awk to add homozygous reference allele calls at every base position. The resulting VCF file was converted to fasta format using vcf2phylip (Ortiz 2019). The resulting fasta file was used as input for maximum likelihood inference using IQtree2 v.2.2.7 (Minh et al. 2020) with the GTR+G+F0 model of molecular evolution and 1000 bootstrap replicates.

### Reconstruction of phylogenomic relationships and gene tree exploration

Prior to gene tree inference, VCF files containing SNP calls for each locus were first converted to fasta format with vcf2phylip (Ortiz 2019). A preliminary ML analysis of a concatenated matrix was conducted using 100 BUSCO genes, with the first five loci from each of the 20 chromosomes concatenated from 570 accessions representing *Glycine* subgenus *Glycine* diploid species diversity plus *G. max* and a phaseoloid legume outgroup (*Amphicarpaea edgeworthii*; Song et al. 2021; NCBI BioProject accession PRJNA663436). The resulting concatenated tree (Supplemental Figure S1) identified the expected major nuclear genome clades of subg. *Glycine* (e.g., Sherman-Broyles et al. 2014a; Landis and Doyle 2023) and allowed selection of 61 accessions (Table 1) representing each known or hypothesized perennial diploid *Glycine* species, with the exception of two I-genome species, *G. aphyonota* and *G. remota* (Landis and Doyle 2023). Two or more accessions were selected to represent each taxon, with the exception of the two other I-genome species, *G. albicans* and *G. lactovirens*, each represented by a single accession. Prior to analyses, alignments for each locus with 61 taxa were summarized with Segul GUI (Chan 2024) to produce locus and taxon stats including missing data and base composition. Because resolution can be lost due to homoplasy created by mixing multiple historical signals (e.g., Schierup and Hein 2000), all loci were tested for evidence of recombination using a subset of taxa representing Dataset 2 (27 individuals) with the PHI analysis in PhiPak (Bruen et al. 2006). In total, 24 out of the 2389 loci displayed evidence of recombination and flagged as problematic in downstream analyses. All loci were used to construct gene trees using IQtree2 v.2.2.7 with the GTR+G+F0 model of molecular evolution and 100 bootstrap replicates. Concatenated trees were also inferred after combining loci together with seqkit v.2.6.1 (Shen et al. 2016a) using the concat function. An approximation of the multi species coalescence was done with ASTRAL v.5.7.8 (Zhang et al. 2018) with the inferred gene trees from IQtree.

Additional subsampling of accessions occurred to investigate phylogenomic differences in the known genome groups of *Glycine.* Specifically, subsets of the A-genome group, B-genome group, and the DEHHaI-genome groups were included. This was done to determine if loci that were informative in Dataset 1 and 2 were also informative for the different genome groups.

Multiple approaches were undertaken to cluster gene trees based on topological similarity to explore gene tree space using Datasets 2 and 3. The Robinson Foulds (RF) distance was calculated in IQTree2, with the resulting distance plotted as a NJ tree. Even after calculating 100 bootstrap replicates, there was not a clear depiction of clusters of gene trees. The R package TreeDist v2.7 (Smith 2022) was then used to cluster gene trees with hierarchical clustering, K-means clustering, and PAM (partition around medoids) after calculating Clustering Info Distance (CID), RF, and Phylogenetic Info Distance (PID) in separate analyses with a maximum number of clusterings between four and 10. Two separate inputs were tested: the inferred gene trees and gene trees with nodes receiving less than 70% bootstrap support collapsed using GoTree v0.4.5 (Lemoine and Gascuel 2021). Loci in different clusters were concatenated with seqkit v.2.6.1 (Shen et al. 2016a) and a concatenated tree was inferred for each. Venn diagrams of loci of interest were plotted using (https://bioinformatics.psb.ugent.be/webtools/Venn/).

Unique topologies were scored from uncollapsed gene trees using TreeTools v1.10.0 (https://github.com/ms609/TreeTools) and phytools v2.1.1 (Revell 2024). Unfortunately, the two approaches did not completely coincide, with TreeTools producing different scores depending on the order of the tip labels. The function “find.unique” in phytools was consistent in scoring unique topologies across all datasets and was used for downstream analyses.

PhyParts v.0.0.1 (https://bitbucket.org/blackrim/phyparts/src/master/) was used to investigate conflict between gene trees and the species tree, concatenated tree, and plastome tree. Cophylo plots highlighting incongruences between trees were created using the R package ape v5.7-1 (Paradis and Schliep 2019). Additional PhyParts analyses were conducted on the three primary clusters recovered from TreeDist using both the concatenated maximum likelihood tree and astral tree. Gene ontology enrichments were done for different subsets of loci based on the PhyParts analyses using the full list of BUSCO genes as the background loci in ShinyGO (Ge et al. 2020) and the *G. max* version 1 genome annotation.

### Divergence time estimation

Divergence times were estimated with the nuclear SNP, mitochondria SNPs, and the plastome trees. Suissa et al. (2024) showed that even though raw branch lengths are drastically different between SNP and whole locus datasets, these differences disappear after divergence time estimation analyses. For all three data sets, we used RelTime (Tamura et al. 2018) as implemented in Mega11 (Tamura et al. 2021). A secondary calibration point of the age of perennial *Glycine* from Zhuang et al. (2022) was used, specifically 6.12 MYA with a normal distribution and standard deviation of 0.435 MYA (5.27-6.97 MYA). For the nuclear dataset, using all loci was computationally challenging based on run time and required RAM. Therefore, only the 1070 loci that showed phylogenetic resolution in the 27-taxon analyses were used. This resulted in a concatenated alignment of 2,406,790 bp. For the plastome data set, the full plastome minus one inverted repeat was used. Given the noticeable differences in branch lengths towards the root of the tree when *A. edgeworthii* or *G. max* were used as outgroups, two analyses were conducted with the 132,532 bp alignment, one with *A. edgeworthii* retained as the outgroup the other with *A. edgeworthii* removed and *G. max* serving as the outgroup. The mitochondria analyses consisted of the 40,175 bp concatenated SNP alignment with *G. max* as the outgroup since mitochondrial SNPs of *A. edgeworthii* were not available. Resulting plots show the 95% confidence interval for dates, and based on Mega plotting, the outgroup is not shown.

### Tests for introgression and network analysis

Recent studies have shown that multiple hybridization/introgression events can mask the signals of each other and make proper identification difficult (Frankel and Ané 2023), therefore multiple different approaches were undertaken. Given that a purely bifurcating tree is likely not the most appropriate representation, especially in Dataset 1 where multiple accessions per species were incorporated, two network approaches were employed. First, a neighbor network was inferred using SplitsTree v4.17.1 (Huson and Bryant 2006) with the concatenated nexus file as input. Additionally, a phylogenetic network was inferred with SNaQ (Solís-Lemus and Ané 2016) implemented in PhyloNetworks v0.11.0 (Solís-Lemus et al. 2017). Due to the computational intensity of PhyloNetworks, only Dataset 3 with seven taxa was used. Zero to four hybridization events were investigated using the ASTRAL tree as the starting tree and the 299 fully resolved gene trees.

An ABBA/BABA test was conducted on Dataset 1 with 61 taxa in D-suite v0.5r47 (Malinsky et al. 2021) and the Dtrios function. The significance threshold was adjusted for multiple testing with a Bonferroni correction of (L/4960) to account for possible trios tested in the data set. To avoid tests involving multiple species of the same genome group, Dataset 3 with the six *Glycine* species was also used with a Bonferroni correction of (L/21). Heat maps using the D-statistic were plotted with ruby (https://github.com/mmatschiner/tutorials/tree/master/analysis_of_introgression_with_snp_data#TestingInSimulations).

### Tests for pseudo-orthology

To assess which of the loci producing topologies incongruent with the species tree were likely to have resulted from pseudo-orthology, we used whole genome assemblies of six *Glycine* species (Zhuang et al. 2022) to identify all BUSCO-matched loci, using BUSCO nucleotide sequences against both the genomic sequence directly, as well as the protein products of the annotated genes. Because *Glycine* experienced a relatively recent whole genome duplication (∼10 MYA; Schmutz et al. 2010), the predominant mechanism for producing pseudo-orthologous loci was hypothesized to be alternate gene loss among homoeologous copies resulting in a mixture of both homoeologues in the gene tree. To define homoeologous loci, we first identified collinear blocks of genes for each pair of species and then estimated the divergence times of putatively homoeologous loci based on the median Ks values of all gene pairs defining those blocks. The next task was to partition the loci corresponding to each BUSCO model into subgroups in such a way that all members of a subgroup were inferred to be orthologous to one another, produced by speciation subsequent to the WGD. For this, we used a single-linkage clustering approach based on the idea that any two loci whose estimated divergence was less than or equal to a maximum value are most likely to have diverged after one of the speciation events that followed the more recent whole genome duplication common to all *Glycine* species; thus, further agglomerative clustering was terminated when there were no more pairs of genes in separate clusters that met the maximal speciation divergence join criterion. We used median Ks=0.08 as the maximum value for speciation-related loci, based on the Ks peaks for between species comparisons which contain modes produced by orthologous gene pairs as well as gene pairs related by the earlier WGD; note that genes matched by BUSCOs but not residing in collinear blocks with other BUSCO-matched loci were excluded from further consideration. The final step in assessing a resulting set of clusters for a given BUSCO was to consider the OTUs assigned to the clusters; essentially, a locus considered as a candidate for pseudo-orthology is one for which there are at least two subgroups produced by the clustering (representing separate homoeologous, non-orthologous groups) and no OTU contributes a gene to more than one of the subgroups. This latter condition implies that one homoeologous copy of a gene has been lost from each of the OTUs, while the former condition implies that the gene loss occurred differentially between the two homoeologous groups.

As an additional test for pseudo-orthology two of the more informative metrics implemented by Frost et al. (2024) for identifying “artifactual orthologs” in taxa lacking genomic sequences or other synteny information, gene tree length and bipartition support, were used with three different data sets. The first two were the gene trees from Dataset 2 and Dataset 3, while the third dataset included the coding sequences from Zhuang et al. (2022) genomes and *G. max* Williams 82 (Schmutz et al. 2010). For this last dataset, the CDS files were first translated to the longest open reading frame using Transdecoder v5.5.0 (https://github.com/TransDecoder/TransDecoder) using the single best prediction flag with no refining of start codons. Resulting files were then used as inputs for Orthofinder v2.5.5 (Emms and Kelly 2019). Fasta files of the identified single copy genes were aligned with MAFFT v7.525 (Katoh and Standley 2013) using the auto function. Gene trees were inferred with IQTree2 v.2.2.7 (Minh et al. 2020) with the GTR+G+F0 model of molecular evolution and 100 bootstrap replicates. Gene trees were rooted with GoTree v0.4.5 (Lemoine and Gascuel 2021) and the species tree was inferred with ASTRAL v.5.7.8 (Zhang et al. 2018) using the rooted gene trees. Tree length and variance of the rooted gene trees were calculated with the get_var_length.py script in SortaDate (Smith et al. 2018). Bipartition support was calculated using the get_bp_genetrees.py script in SortaDate with the rooted gene trees and the ASTRAL tree used as inputs. Tree length and bipartition support were plotted in R using strip plots with the average values calculated and used as the filtering thresholds for both parameters as done by (Frost et al. 2024). Loci that were outliers for both tree length bipartition support were designated as problematic loci for pseudo-orthology.

## Results

### Species and concatenated trees for subgenus Glycine

The topologies of the concatenated tree and the ASTRAL species tree for the BUSCO genes for 61 perennial *Glycine* accessions and outgroups (Dataset 1) were congruent (Figure 1; Supplemental Figure S2). All of the known nuclear genome groups of the subgenus were identified with strong bootstrap support, as were relationships among genome groups, including the sister relationship of *G. falcata* (F-genome) to the remaining perennial species clades; the split of remaining groups into B+C genomes vs. a clade comprising taxa from the A-, D-, E-, H-, Ha- and I-genomes (subsequently called the “ADEHI clade”); the resolution of the latter clade as A(I(DEH/Ha)); and the sister relationships of the D and E and the H and Ha genomes. Many of the deeper nodes have full bootstrap support (100%), however the normalized quartet scores from ASTRAL, which show the percentage of quartets in the gene trees that agree with the marked branch, are quite low (Figure 1). With the low resolution of many of the individual loci (see below), this low quartet score when taking into account all loci is not unexpected. Resolution of *G. canescens* and *G. clandestina*, which are two closely related species complexes that come out as paraphyletic, and other issues such as those involving informally named taxa in the B-, H-, and Ha-genomes, require further study, and will be addressed using focused studies of the full >500 accession dataset, drawing on the exploration of gene tree topology space for the BUSCO genes here. Although all nodes were strongly supported in both the ASTRAL and concatenation trees, PhyParts analysis of these species trees showed that for some nodes, many (in some cases the majority) gene trees conflicted with the species tree (Figure 2) and the concatenated tree (Supplemental Figure S3).

**Figure 1.**
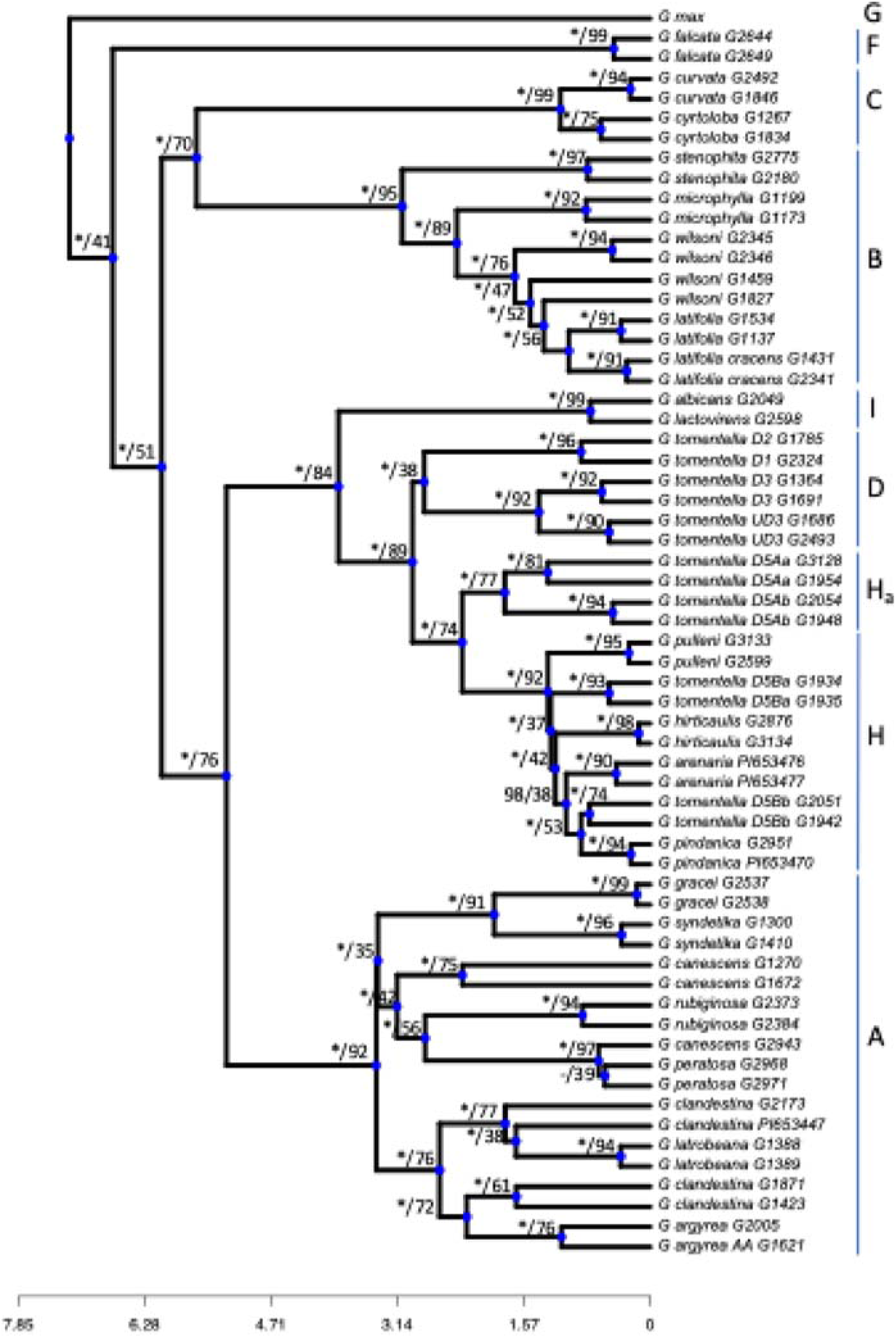
ASTRAL divergence time and support rooted with *Amphicarpaea edgeworthii* (not shown). Bootstrap values from the concatenated maximum likelihood inference and the normalized quartet support scores (option “-t 1”) from the ASTRAL analyses are shown at each node. Nodes with 100% bootstrap support are shown with *. The single node not resolved in the maximum likelihood concatenated tree is indicated with “-”

**Supplemental Figure S2.**
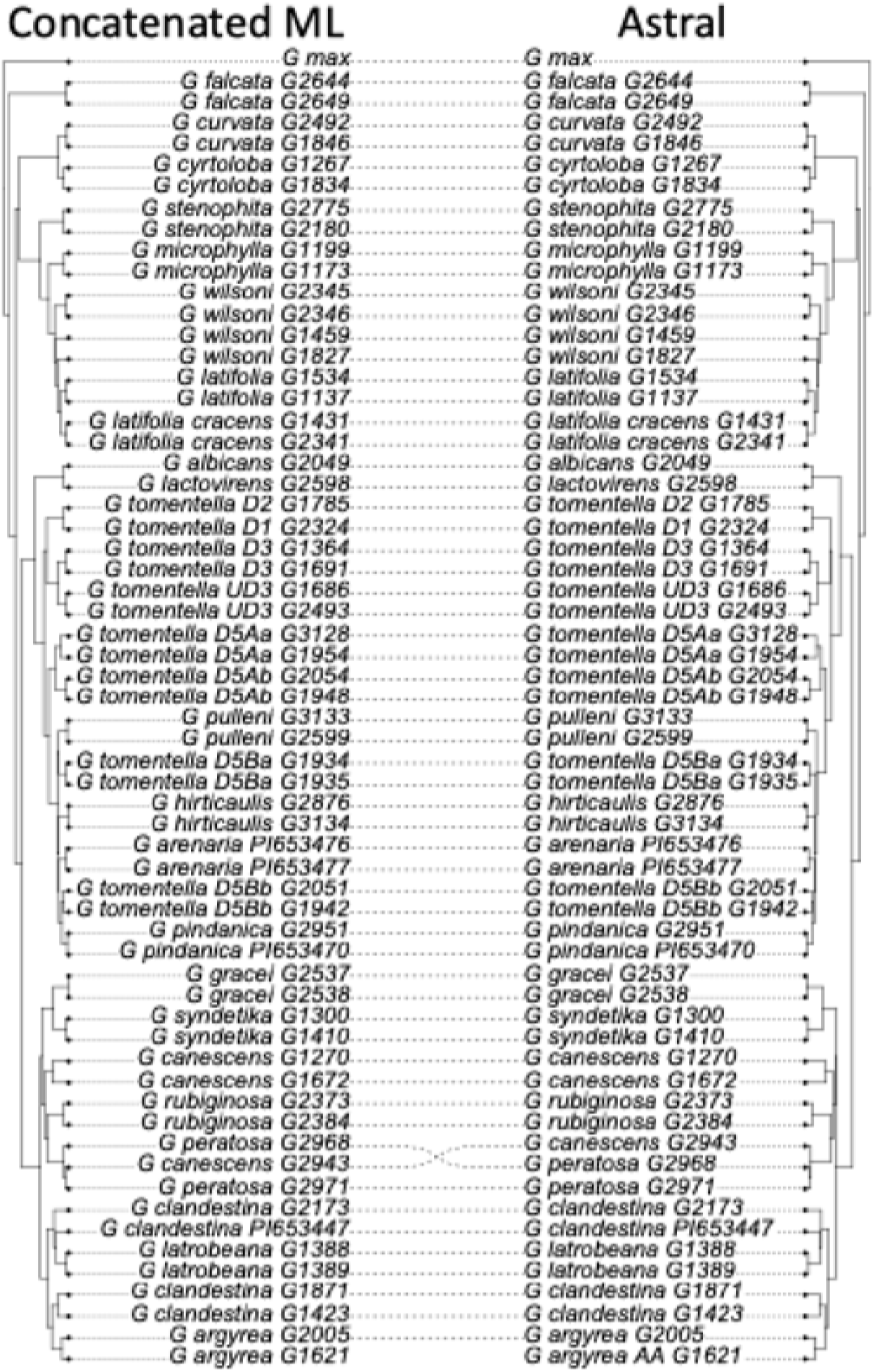
Cophylo plot between Astral and concatenated trees for the 61-taxon dataset.

**Figure 2.**
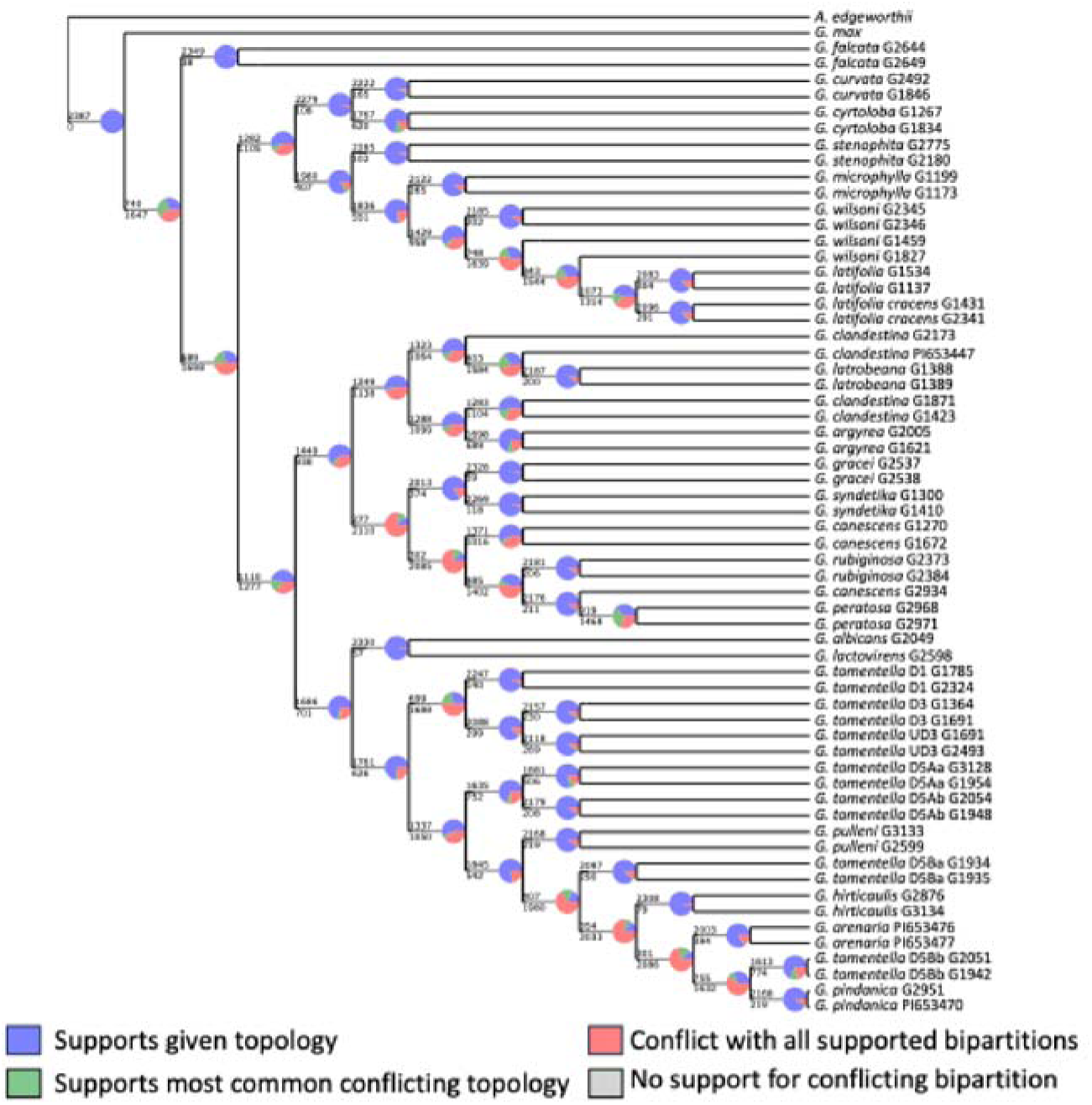
PhyParts showing ASTRAL topology and individual gene tree support.

**Supplemental Figure S3.**
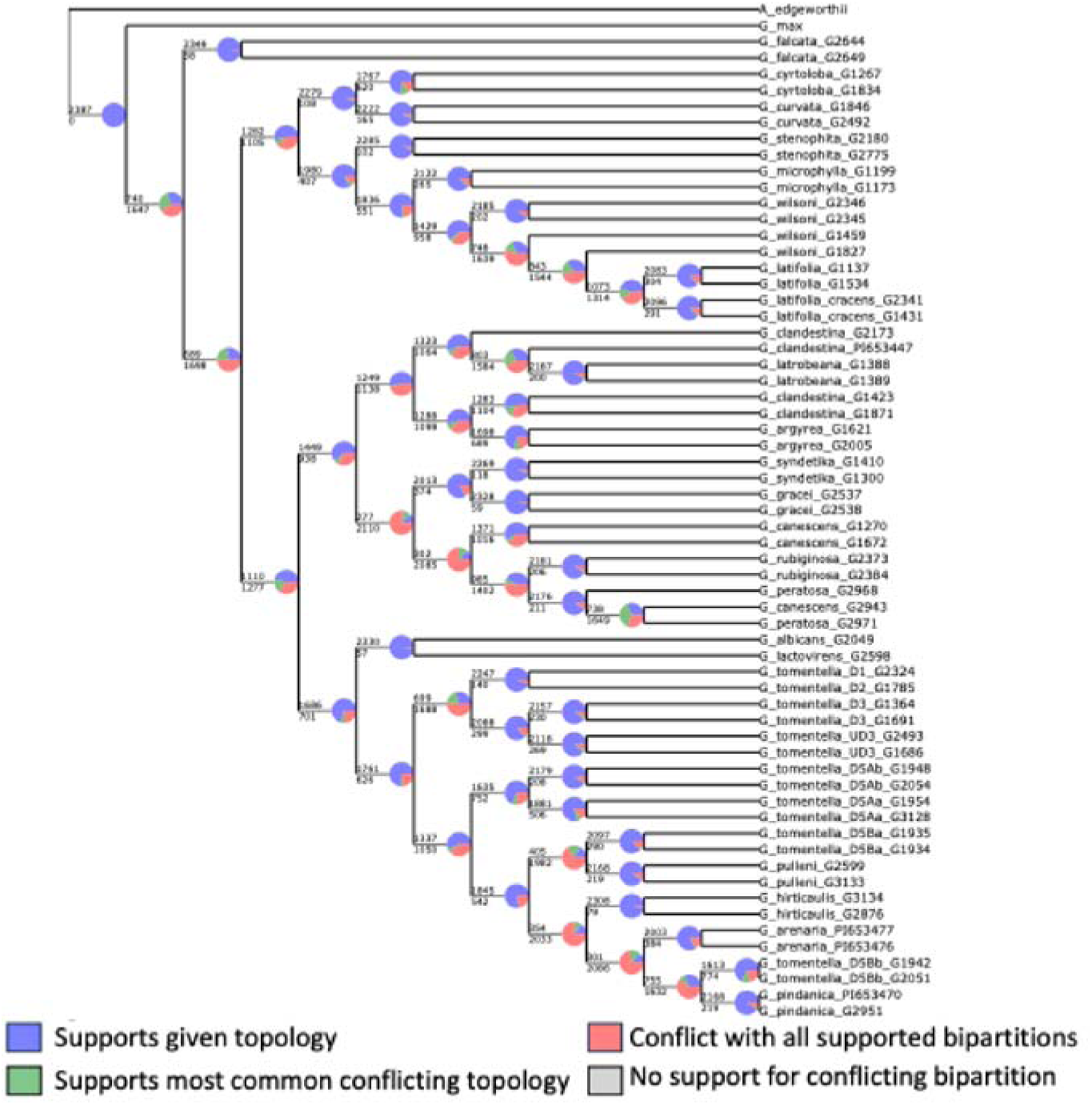
PhyParts showing concatenated topology and individual gene tree support.

### Incongruence involving the plastid and mitochondrial genome

ML analysis of the plastid genome identified a plastome tree that was well-resolved, with 100% bootstrap support for all but three nodes (Supplemental Figure S4). As was expected based on earlier studies using chloroplast RFLPs and complete plastid genome sequences from a small number of species, the overall organellar genome topology was highly incongruent with the species tree (Figure 3). Notably, *G. falcata* plastomes are members of an A-plastome clade, which comprises plastid genomes from all accessions sampled from species of the ADEHI clade. The *G. falcata* plastome clade is sister to a clade comprising all ADEHI clade plastomes with the exception of both accessions of *G. rubiginosa* plus one of the four sampled accessions of *G. clandestina*. This “rubiginosa” clade is relatively weakly supported (76% bootstrap) as sister to the remaining A-plastomes, and thus also highly incongruent with the species tree, where they and all *G. clandestina* accessions are placed robustly in the A-genome group. Other aspects of the A-plastome topology also disagree with the species tree. The A-plastome clade is sister to a B+C-plastome clade, both of whose sister clades correspond to the B- and C-genome groupings as in the species tree. However, as with the A-plastome group, the topology within the B-plastome clade does not track B-genome species relationships (Figure 1, Supplemental Figure S2).

**Figure 3.**
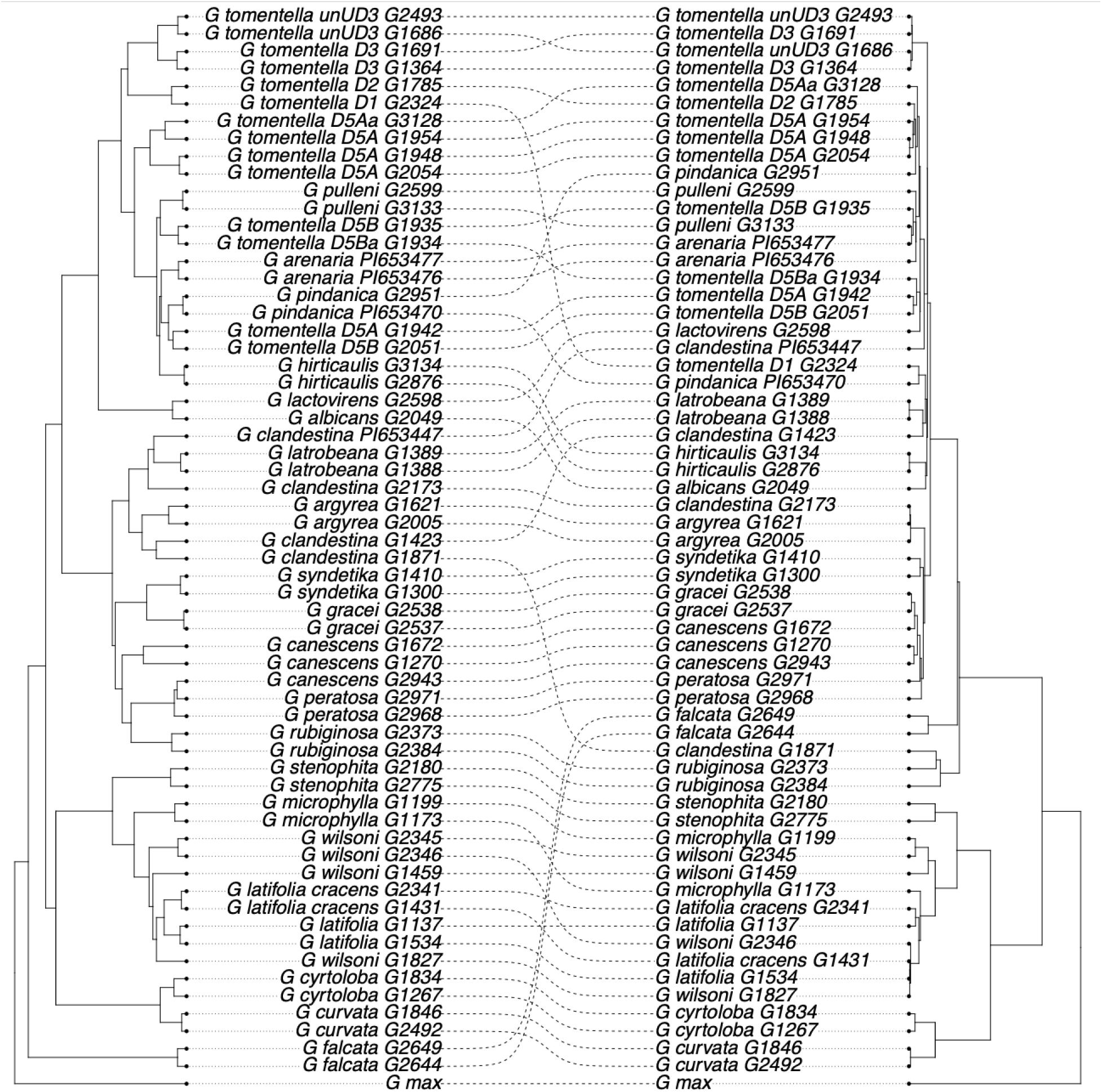
Cophylo plot between nuclear (left) and plastome (right) trees. [JJD will add highlighting to show contrasting positions of G. falcata]

**Supplemental Figure S4.**
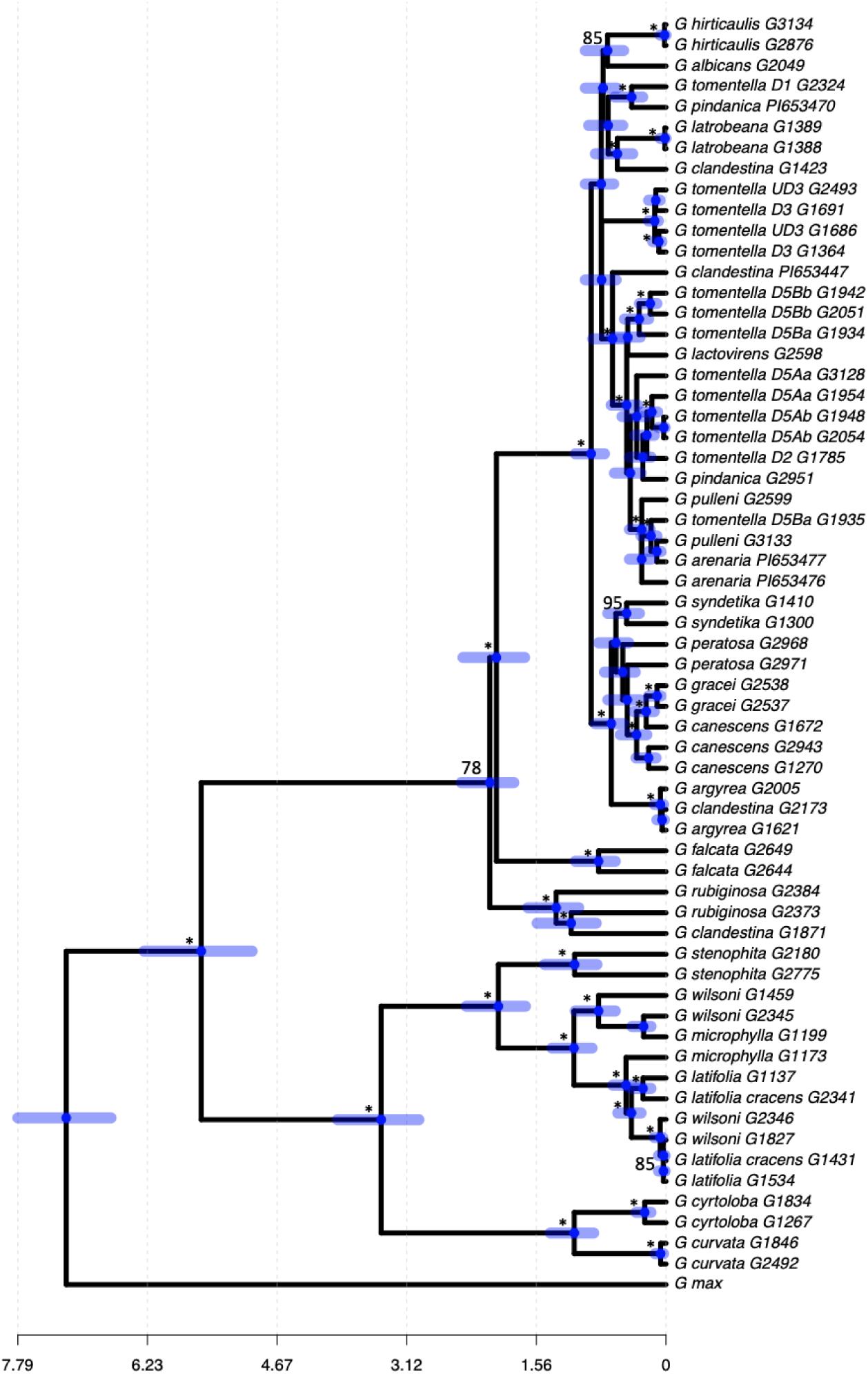
Dated plastome tree rooted with Amphicarpaea edgeworthii (not shown). Support values above 70% are shown, and nodes having 100% bootstrap support are labeled with *.

The ML concatenated SNP mitochondrial tree was fully resolved but more weakly supported than the plastome tree: 28 of 55 nodes had 100% bootstrap support, and all but five of the remaining 27 nodes had >70% support (Supplemental Figure S5). The two cytoplasmic genomes share several key features (Supplemental Figure S6), notably the dichotomy between a B+C genome clade and the remainder of the perennials, with G. falcata belonging to the latter group rather than being sister to all perennials as in the species tree (Figure 1, Supplemental Figure S2). Unlike the plastome tree, however, a *G. falcata* mitogenome clade was sister to all remaining ADEH/HaI plastomes, switching places with a “rubiginosa” mitogenome clade relative to the plastome tree (Supplemental Figure S6). In both the plastome and mitogenome trees, however, these relationships received less than 100% support. The mitogenome and plastome trees disagreed topologically in other relationships, including the membership of their respective, fully supported “rubiginosa” clades. In both trees, the two *G. rubiginosa* accessions were joined by the same *G. clandestina* accession; however, the mitogenome clade included a second *G. clandestina* accession, whose plastome grouped strongly elsewhere in the plastome tree (Supplemental Figure S6).

**Supplemental Figure S5.**
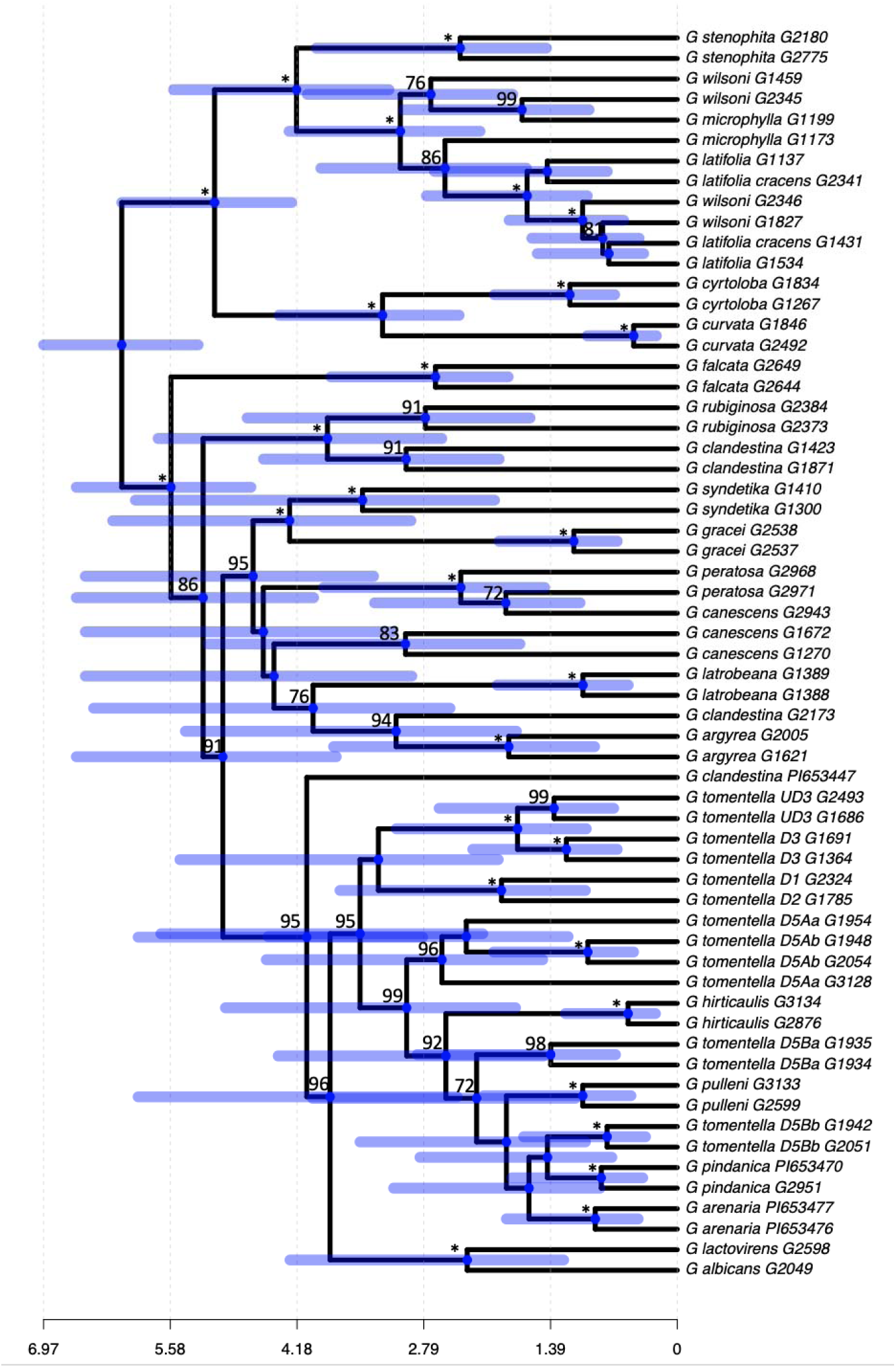
Dated Mitochondrial tree rooted with G. max (not shown). Nodes with bootstrap support above 70% are shown, with nodes with 100% support indicated by *.

**Supplemental Figure S6.**
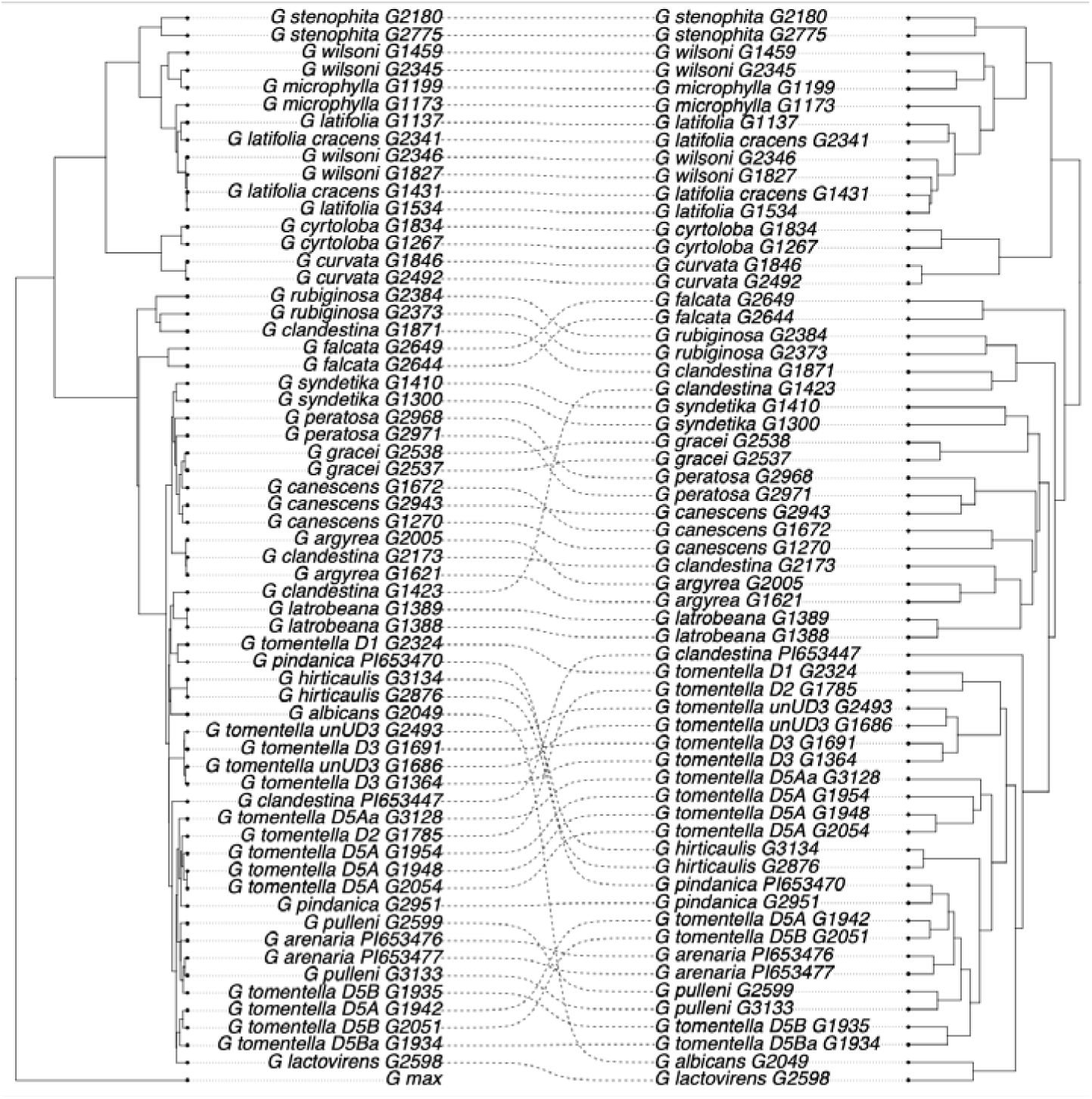
Cophylo plot of plastome v mitochondrial trees

The placement of G. falcata organellar genomes within A-plastome and A-mitogenome clades rather than as sister to all other perennial Glycine genomes suggests that the incongruence observed with the species tree is likely due to introgression rather than to deep coalescence. The likelihood that the G. falcata plastome was obtained by introgression from an A-plastome/mitogenome taxon raises the question of whether there is a legacy of this event in the nuclear genome, specifically among the BUSCO genes used to construct the species tree. A PhyParts analysis was conducted on a tree with the plastome topology and showed that 144 of these nuclear loci placed G. falcata with ADEHI taxa, as in the plastome and mitogenome trees, rather than as sister to all perennial taxa (Supplemental Figure S7). Enrichment analyses for gene ontology terms did not identify any statistically significant enrichment in these 144 loci, but among them were numerous loci with cytoplasmic functions, such as 15 genes involved with chloroplast function and seven genes associated with the mitochondria (23 out of 144 or 16% associated with cytoplasmic functions) (Supplemental Table S1).

**Supplemental Figure S7.**
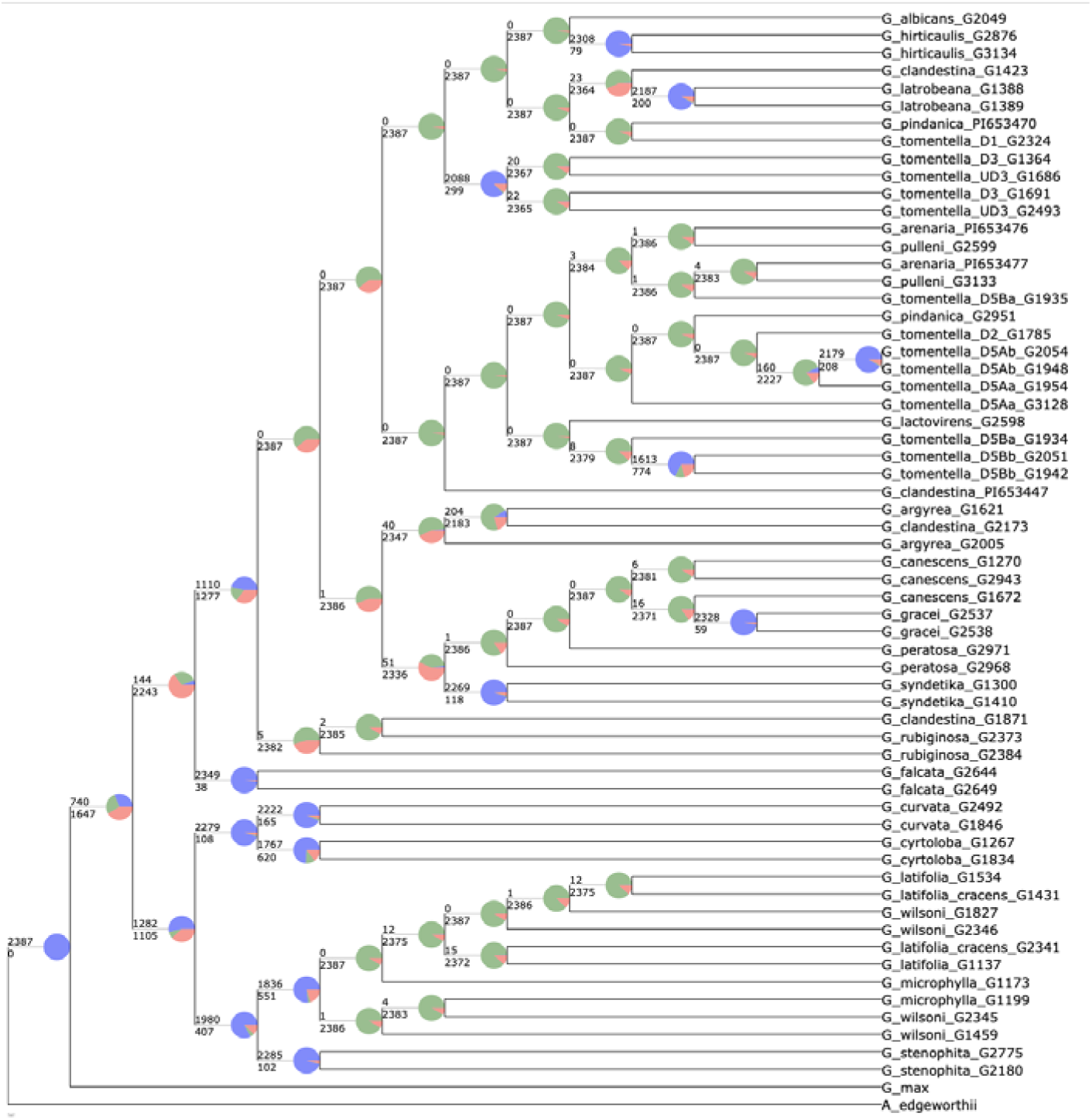
PhyParts of individual BUSCO genes to the plastome phylogeny

### Gene tree topology space is flat for all but the smallest dataset

Using phytools we determined the number of unique topologies found among the loci used in the 61-taxon analysis, with the goal of identifying loci tracking the species tree, nuclear loci introgressed along with the cytoplasm into G. falcata, loci showing ILS, and other histories potentially of biological interest in gene tree topology space. We found that all BUSCO loci had unique topologies. When including the concatenated and ASTRAL trees into the analysis, no single locus was entirely congruent with either the concatenated or the ASTRAL tree. Reduction of the dataset to 27 taxa comprising single accessions for each species plus the two outgroups for all nuclear loci and the plastome (Dataset 2; Table 1) also showed no complete congruence with any locus, except when trees with nodes with less than 70% bootstrap support were collapsed, in which case only a few instances of loci with shared topologies were seen, and these involved highly unresolved trees with only one or two pairs of taxa forming clades.

We reduced the data set to include only six *Glycine* taxa plus the *Amphicarpaea* outgroup (Dataset 3; Table 1). The ML analysis of the BUSCO loci for these taxa identified 299 loci whose gene trees were fully resolved and with >=70% bootstrap support for all nodes (Supplemental Table S2). Concatenation and ASTRAL trees were constructed from these 299 loci (Supplemental Figure S8); phytools identified 36 different unrooted topologies among these 299 loci, out of the possible 105 resolved bifurcating topologies, and 54 of 945 possible rooted topologies. The most common unrooted topology was found in 186 loci; 141 of these trees had the correctly rooted species tree topology, with the remaining 45 having six alternative rootings (Figure 4, Supplemental Figure S9). The second most common unrooted topology, shared by 25 loci, included 15 loci with the correctly-rooted plastome topology and 10 loci with five other rooted topologies.

**Figure 4.**
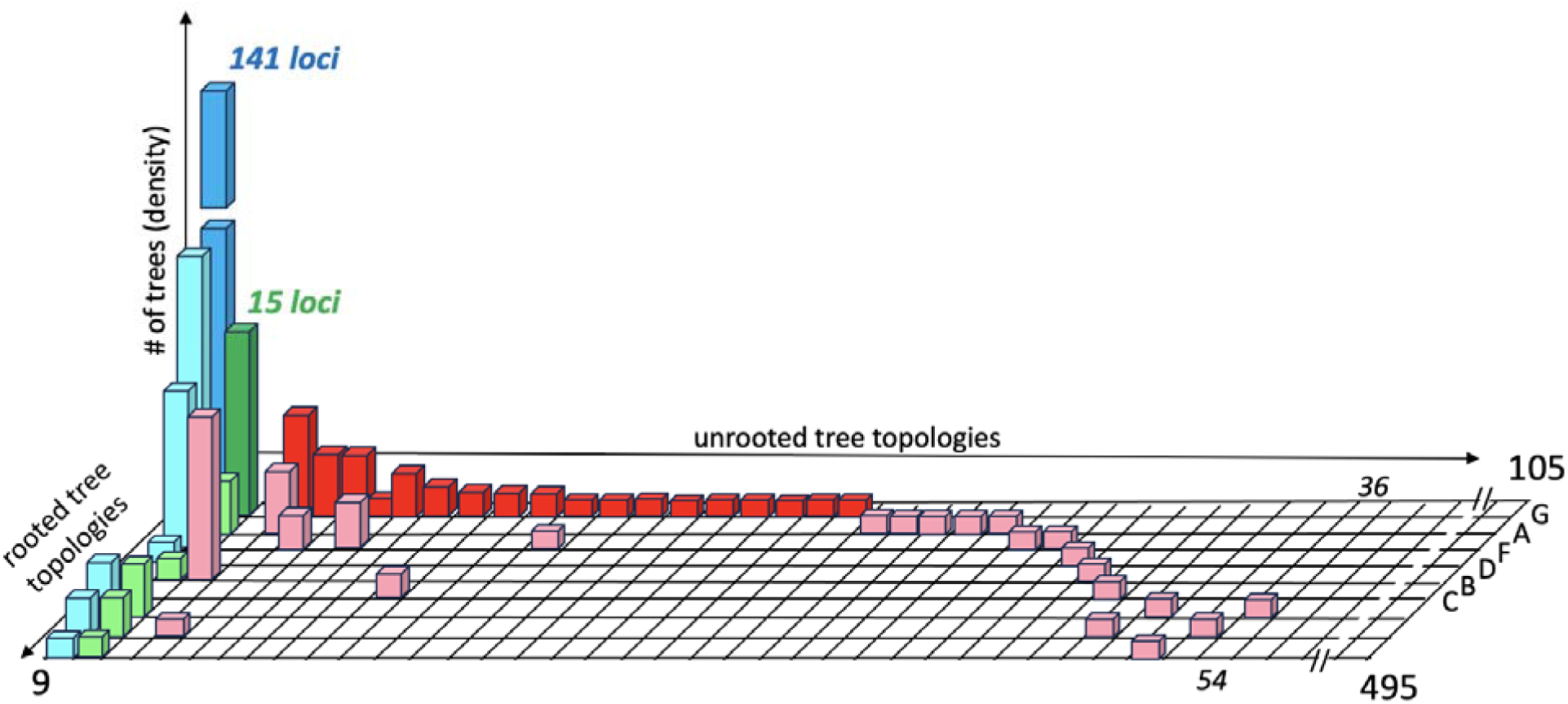
Occupancy of gene tree topology space for the Glycine six ingroup taxon dataset. There are 105 unrooted bifurcating, resolved topologies, with nine possible rootings for each unrooted tree, for a total of 495 possible rooted topologies (squares on checkerboard, numbered from 1 at the top left corner to 495 at the bottom right corner).

**Supplemental Figure S8.**
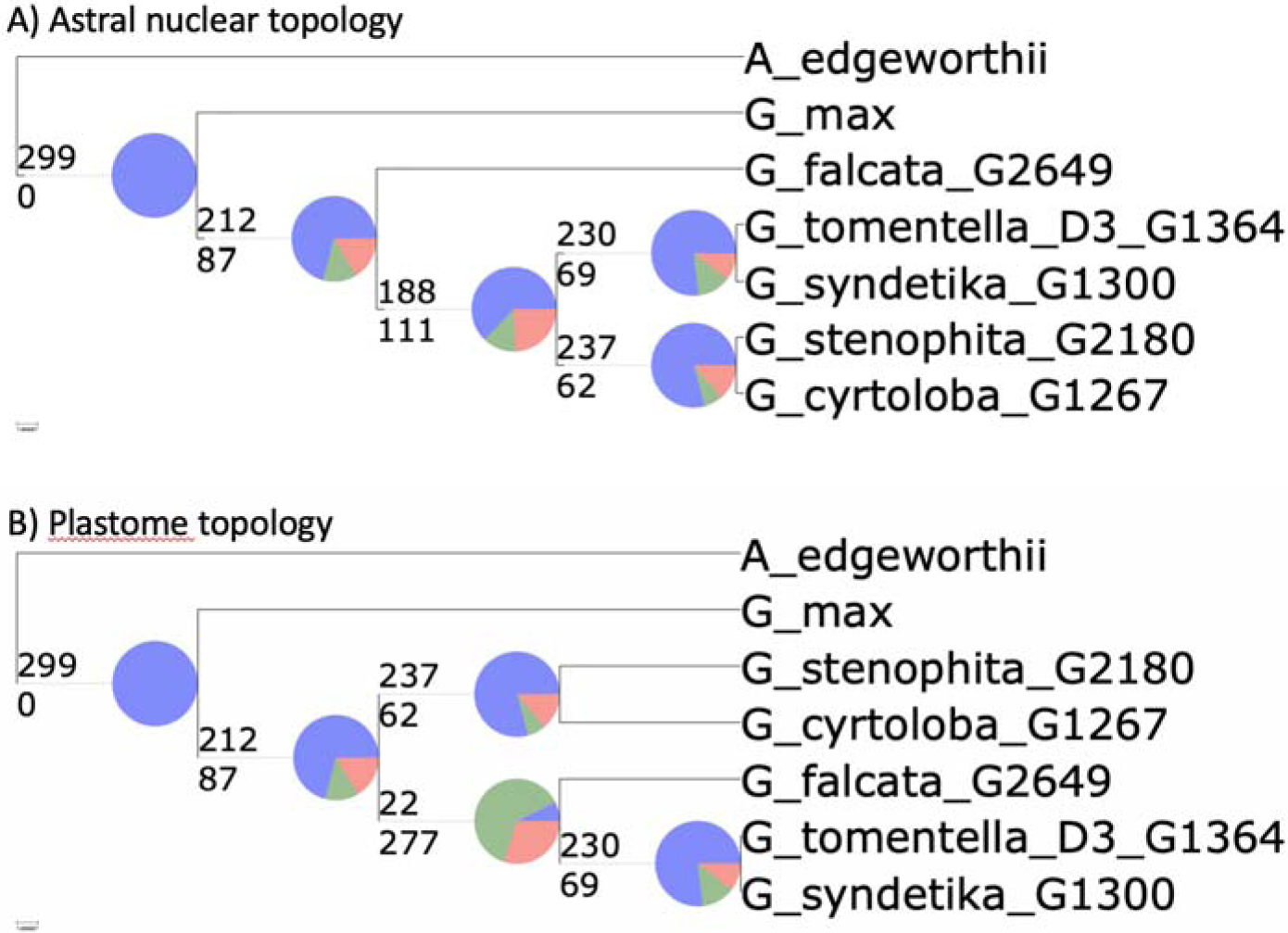
ASTRAL trees and plastome with PhyParts for Dataset 3 representing the taxon sampling of Zhuang et al. constructed from fully resolved gene trees of 299 loci.

**Supplemental Figure S9.**
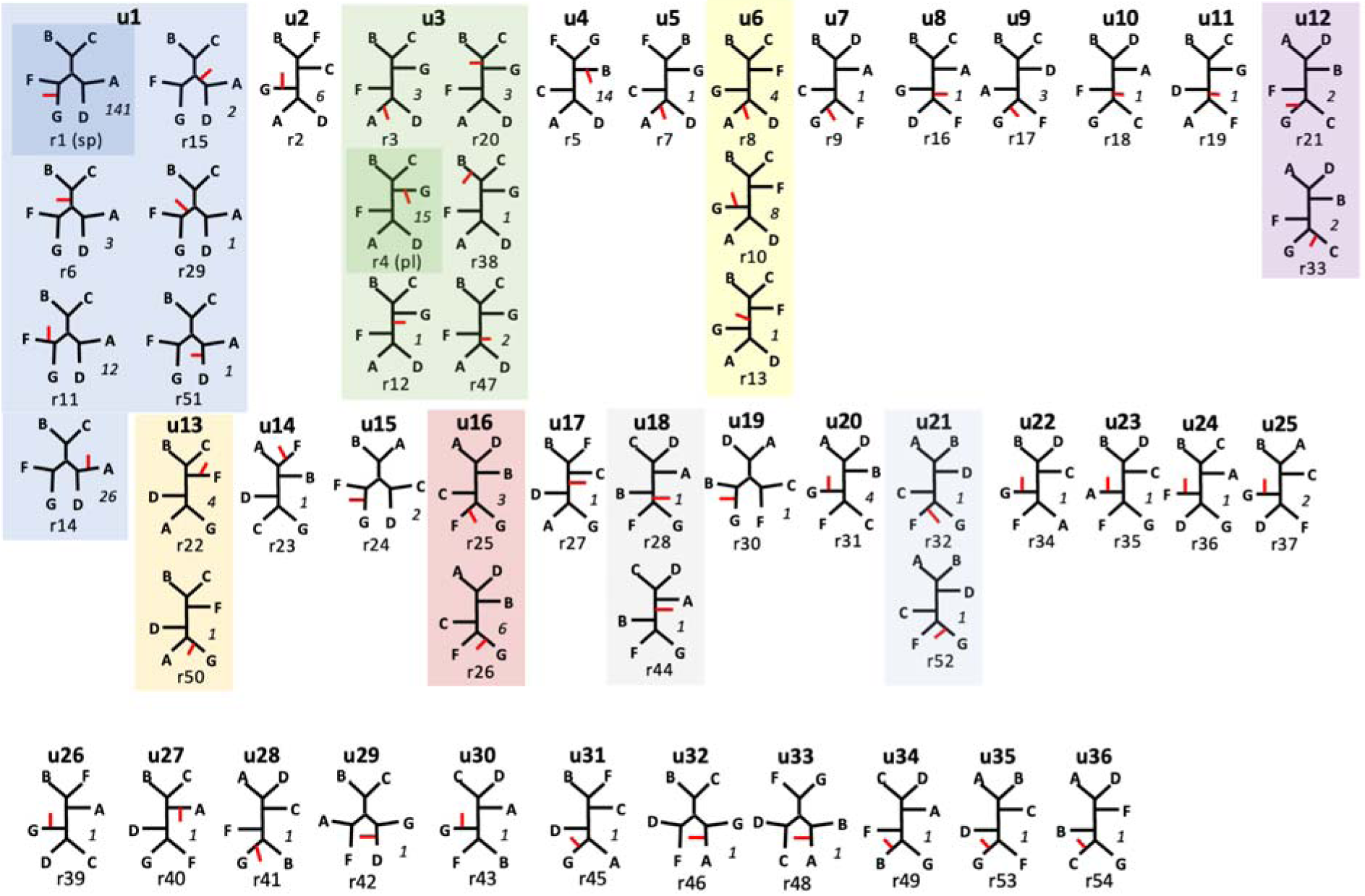
Unrooted (u1-u36) and rooted (r1-r54) topologies for 299 fully resolved and robustly supported gene trees from Dataset 3. Taxa are labeled by genome group. Position of root is shown by a red line. The number of loci with each rooted topology is shown to the right of each tree in italics. Unrooted topologies represented by more than one rooting are colored. Tree r1 (“sp”, dark blue background) is the species tree; tree r4 (“pl”, dark green background) is the plastome tree.

Thus, gene tree topology space for this small dataset is not flat, but is dominated by the species tree (47% of the 299 loci with fully resolved and correctly rooted trees), followed by the correctly rooted plastome topology (5% of the 299 loci; 9.5% of loci with a rooted topology other than that of the species tree). These numbers increase considerably if unrooted topologies consistent with the species tree (186 loci, 62% of all loci) or the plastome topology (25 loci, 12.8%) are considered.

Incorrect rooting was a major contributor to populating gene tree topology space, with 100 loci (33.4%) being incorrectly rooted. This represented 63.3% of the 158 loci that did not track the rooted species tree, and 69.9% of resolved loci having neither the rooted species tree or rooted plastome topology. Given the long distance from the outgroup to the *Glycine* MRCA and the short internodes at the base of the *Glycine* phylogeny (Figure 1, Supplemental Figure S2), unequal distribution of missing data, and the lower quality of the *Amphicarpaea* genome assembly, rooting problems are expected (Supplemental Table S2, Supplemental Table S3). Indeed, 22 of the incorrectly rooted loci were rooted at *G. falcata*, the only member of the clade sister to the remaining perennial species and thus a long branch; of these, 12 were incorrect rootings of the species tree topology. An additional 34 loci were rooted at the A-genome representative, *G. syndetika*, including 26 loci with the unrooted species tree topology. For most of these loci, this rooting appears to be an artifact of using another A-genome species, *G. canescens*, as the reference.

In total, 18 different topologies other than the species tree or the plastome topology were found among the 56 loci with correctly rooted trees, 10 of which were unique to a single locus and the remainder in eight or fewer loci (Figure 4). Two of the most frequent of these topologies, found with 14 and six loci, are closely related to one another and can be explained as a failure to resolve the B- and C-genome representatives as a clade and placing them in the two alternative positions relative to (A,D) and (G,F) clades. The other topology represented by six loci has B sister to F, and can be explained by long branch attraction. More interesting is the rooted topology that is the counterpart of the plastome tree, with F resolved as sister to (B,C) instead of (A,D). This topology is represented by eight loci; an additional five loci have two alternative rootings of the same unrooted topology. The frequency of this topology could represent lineage sorting at the base of the perennial *Glycine* phylogeny. Although this would not account for the shallow coalescence of the *G. falcata* plastome as part of the A-plastome group, lineage sorting could weaken the hypothesis that nuclear loci introgressed into *G. falcata* along with the plastome.

### Trees are not clustered meaningfully in gene tree topology space

The phytools analyses showed that for all but the smallest dataset, gene tree topology space does not contain peaks populated by the trees from more than one locus. However, phytools does not provide information on the relative positions of topologies in tree space. It is possible that, in the larger datasets, related topologies, though each unique, are clustered in a biologically meaningful way, for example with variants of the species tree topology occurring in proximity to one another. To test this hypothesis, we used the more computationally tractable 27-taxon Dataset 2 (Table 1) by selecting representative taxa from each major clade. Concatenation and ASTRAL analyses of this dataset produced similar topologies (Supplemental Figure S10), which included groupings similar to those found with Datasets 3 and 4 (Figure 1, Supplemental Figure S2, Supplemental Figure S8).

First, we calculated pairwise Robinson-Foulds (RF) distances among all BUSCO gene trees for Dataset 2, plus topologies representing the species tree and plastome trees, and estimated relationships using Neighbor-Joining which produced a complex tree with mostly short internal branches whose interpretation was not obvious. The plastome and species tree topologies were separated from one another, but embedded among trees that bore no obvious differentiating characteristics (Supplemental Figure S11).

Silhouette coefficient analysis of several clustering methods on the BUSCO gene trees (see Methods) supported the existence of two clusters, each consisting of trees for over 1,000 loci (Supplemental Figure S12). One cluster (Cluster 1 with 1,359 loci) contained trees averaging ∼75% (19.4/26) of nodes resolved, whereas trees in Cluster 2 (1,027 loci) were less resolved (average 46%; 12.0/26). Cluster 1 included both the species tree and the plastome tree, both of which are fully resolved. Cluster 1 included 52 loci showing statistically significant levels of recombination, with eight remaining after multiple-test correction. Cluster 2 included 120 such loci, with 16 remaining after Bonferroni correction. In both clusters, recombinant trees had slightly lower average numbers of supported nodes than trees with no evidence of recombination (Cluster 1: 18.5 for 52 recombinant loci vs. 19.40 for nonrecombinant loci; Cluster 2: 12.04 for 120 recombinant loci vs. 11.57 for nonrecombinant loci). Therefore, the presence of recombination was not sufficient to account for the 7.4 node difference in average resolution between loci belonging to the two different clusters.

**Supplemental Figure S10.**
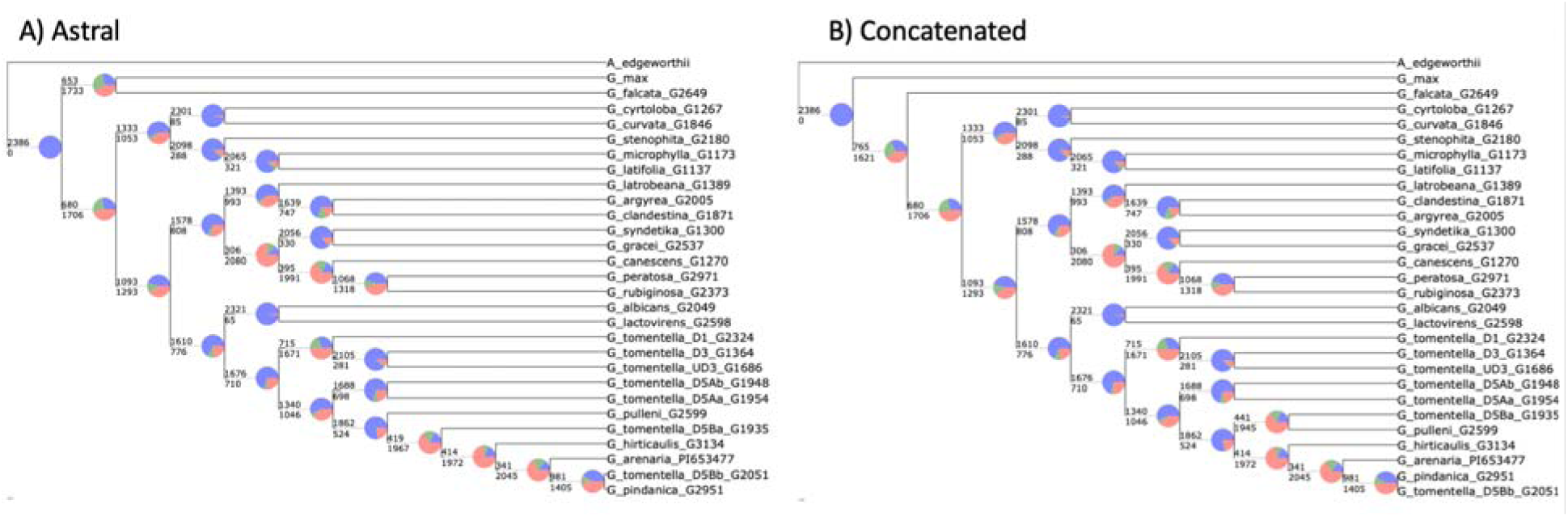
Concatenation and ASTRAL trees with PhyParts for 2386 Dataset 2 loci.

**Supplemental Figure S11.**
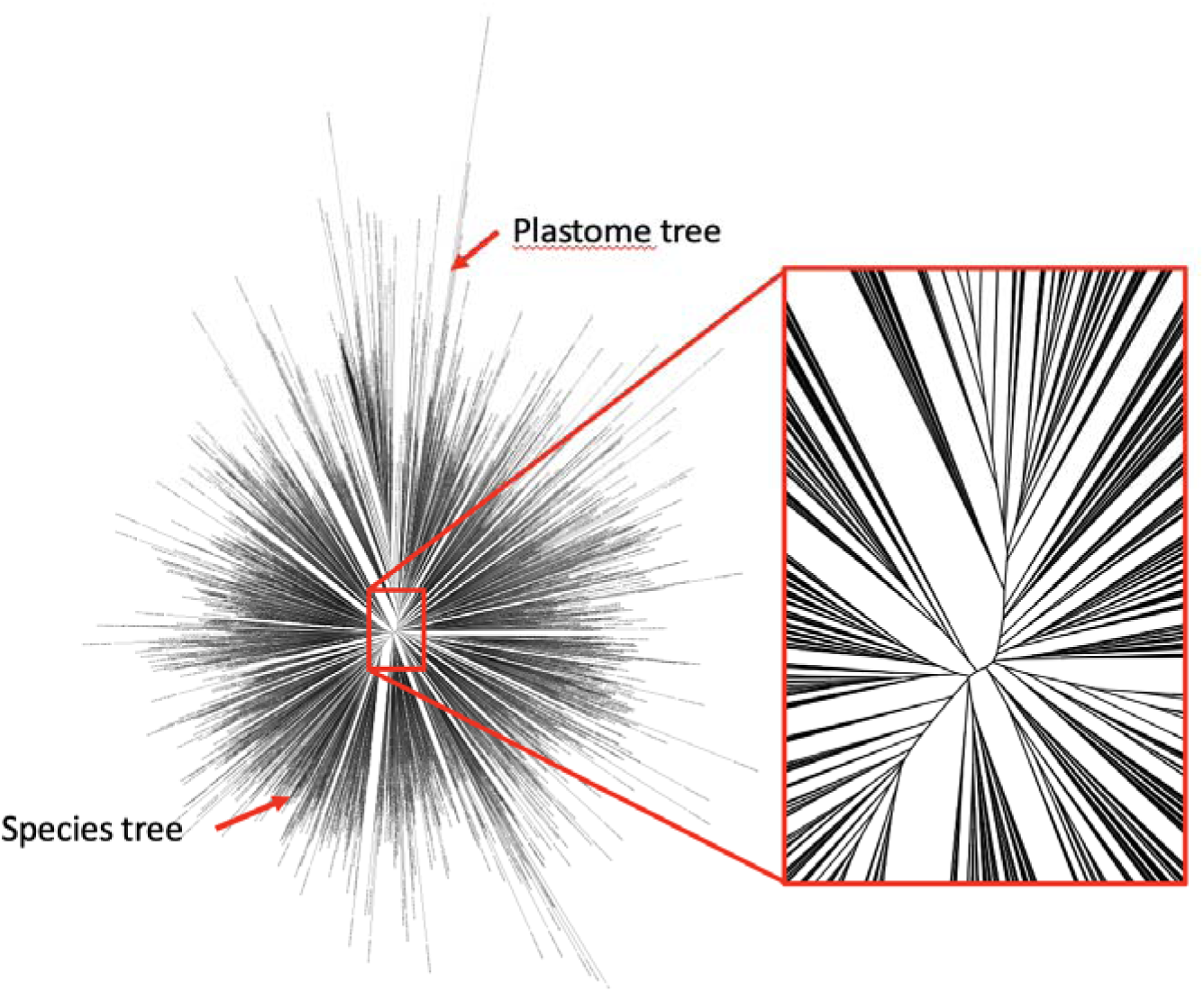
Neighbor-joining tree of pairwise RF distances between 2386 gene trees for Dataset 2, plus species tree and plastome topologies.

**Supplemental Figure S12:**
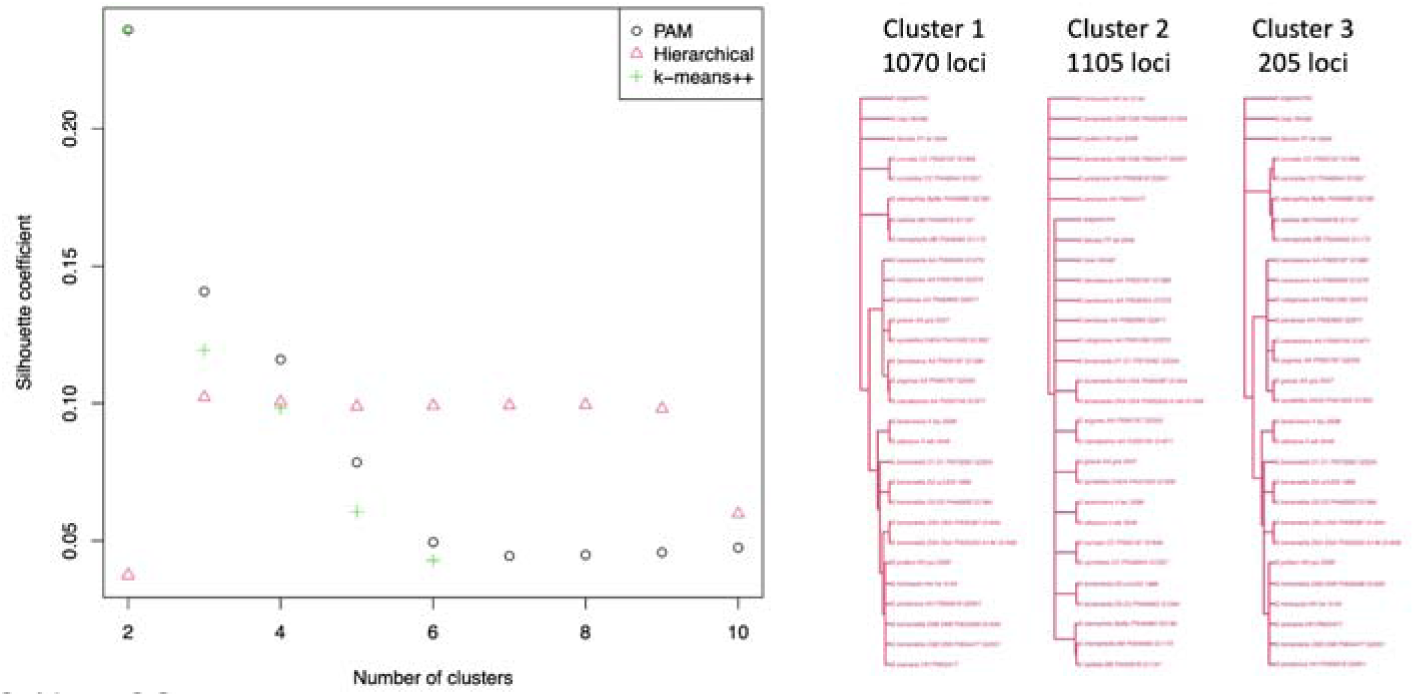
Silhouette coefficient analysis for choice of cluster number in gene tree space exploration and the resulting consensus trees when using three clusters based on the hierarchical clustering metric.

Loci in Cluster 2 may possibly be evolving too slowly to resolve relationships within Glycine, whose species have radiated within the last 6-7 MY. However, median branch length (including outgroups) was virtually identical in the two clusters (Cluster 1: 0.0081; Cluster 2: 0.0084), and Cluster 2 actually had a considerably higher average branch length than Cluster 1, but not a significantly higher variance, suggesting that the difference was not due to the presence of a few long branches (Cluster 1: 0.0227; Cluster : 0.0759).

Species trees and concatenated trees were constructed separately for the two clusters. Concatenation trees (Supplemental Figure S13, Supplemental Figure S14) included the same species relationships obtained for Dataset 1 (Figure 1). Although the Cluster 1 ASTRAL tree was also identical to the species tree from the entire dataset, the ASTRAL tree from Cluster 2 was less resolved and differed from the species tree in placing *G. falcata* sister to the (B, C) clade, albeit with weak support (Supplemental Figure S13, Supplemental Figure S14). From these results we conclude that the two Clusters do not differ in their overall phylogenetic signal–both sets of loci track the same species tree. However, for the generally less resolved trees of Cluster 2, the species tree signal is more effectively obtained by concatenation than by using a two-step coalescent species tree approach.

**Supplemental Figure S13:**
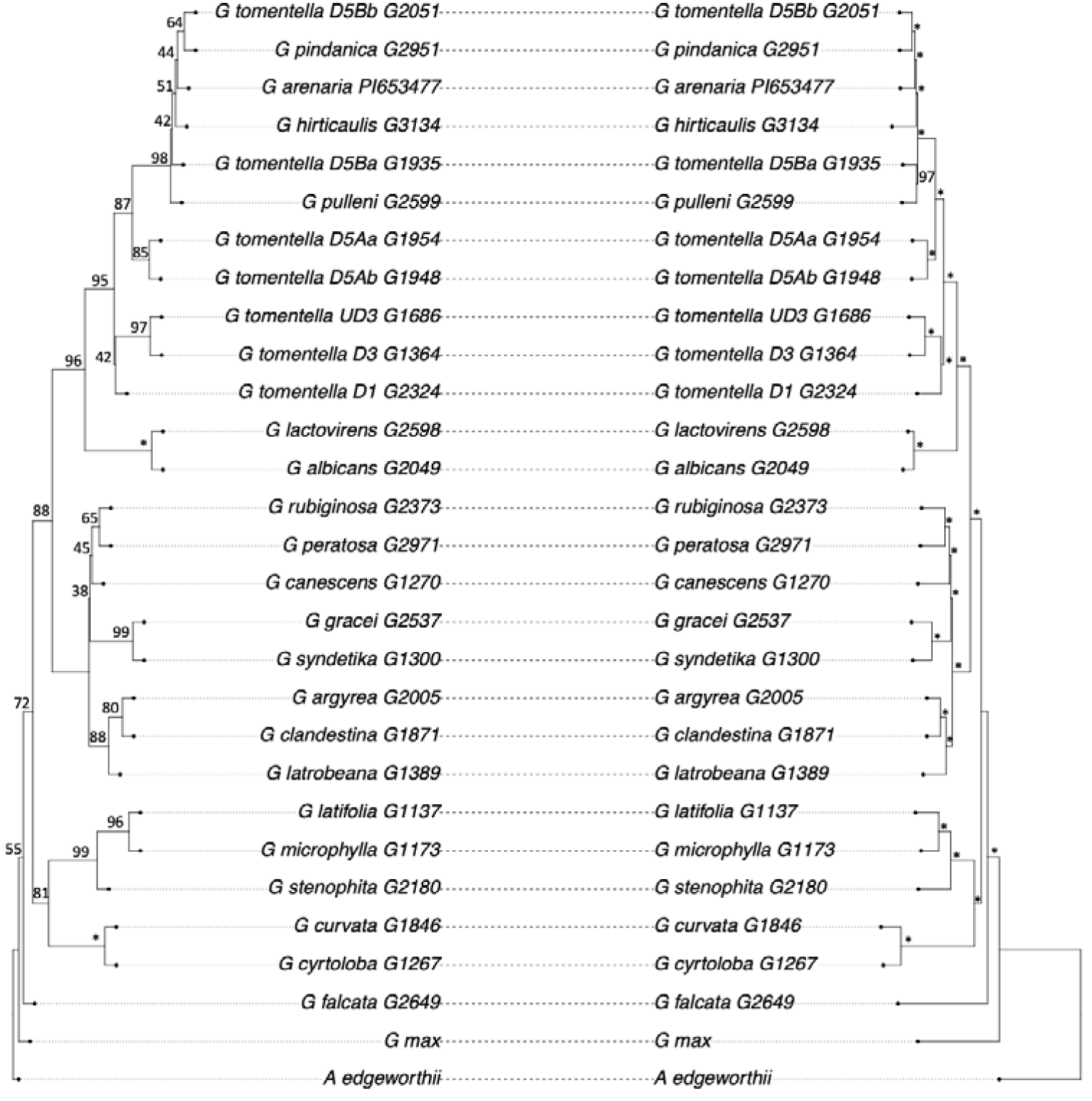
Cluster 1 cophylo ASTRAL and concatenation. Support values (quartet 1 score for ASTRAL and bootstrap for concatenation) are shown. * denotes 100% support.

**Supplemental Figure S14:**
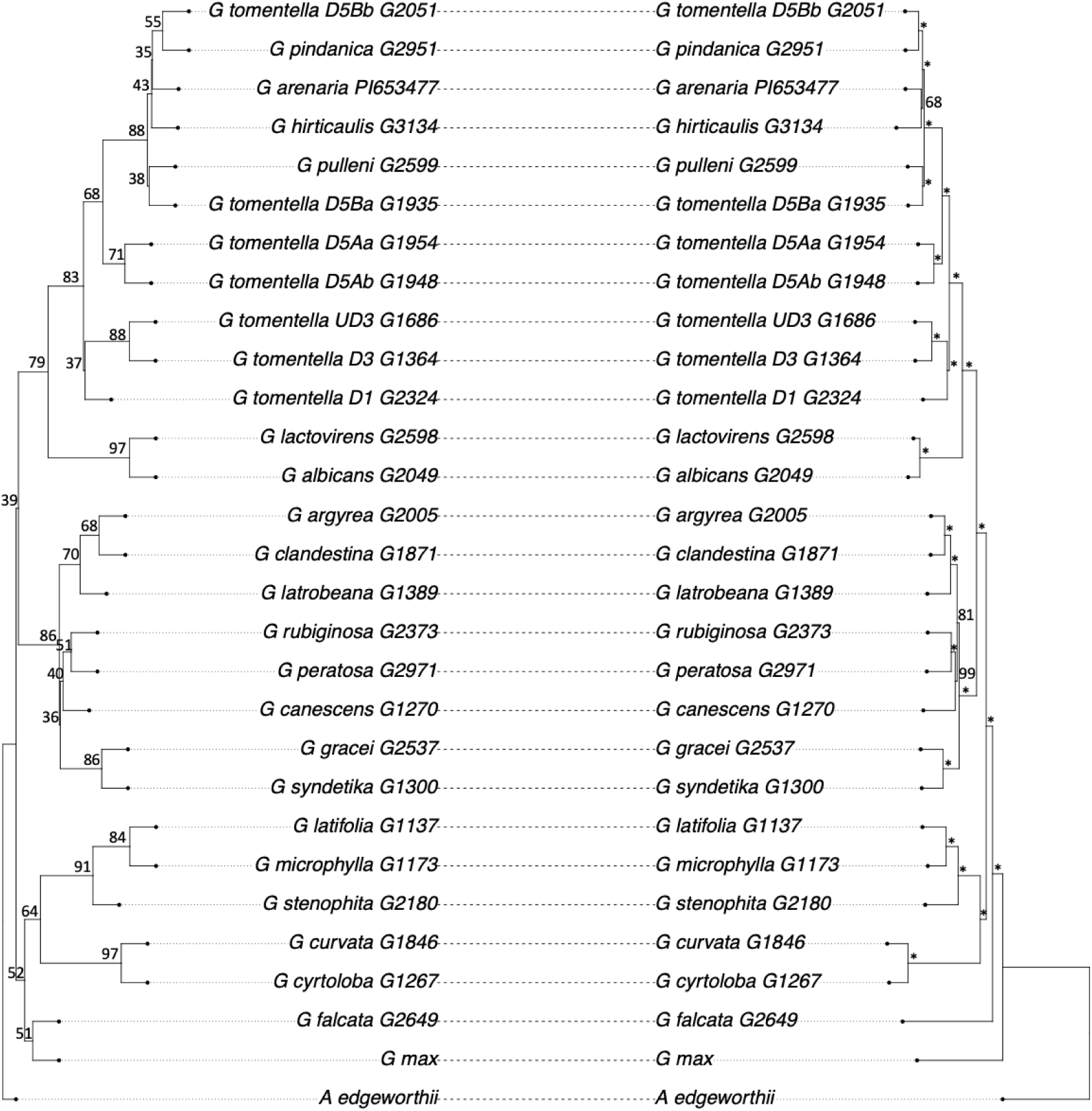
Cluster 2 cophylo ASTRAL and concatenation. Support values (quartet 1 score for ASTRAL and bootstrap for concatenation) are shown. * denotes 100% support.

**Supplemental Figure S15:**
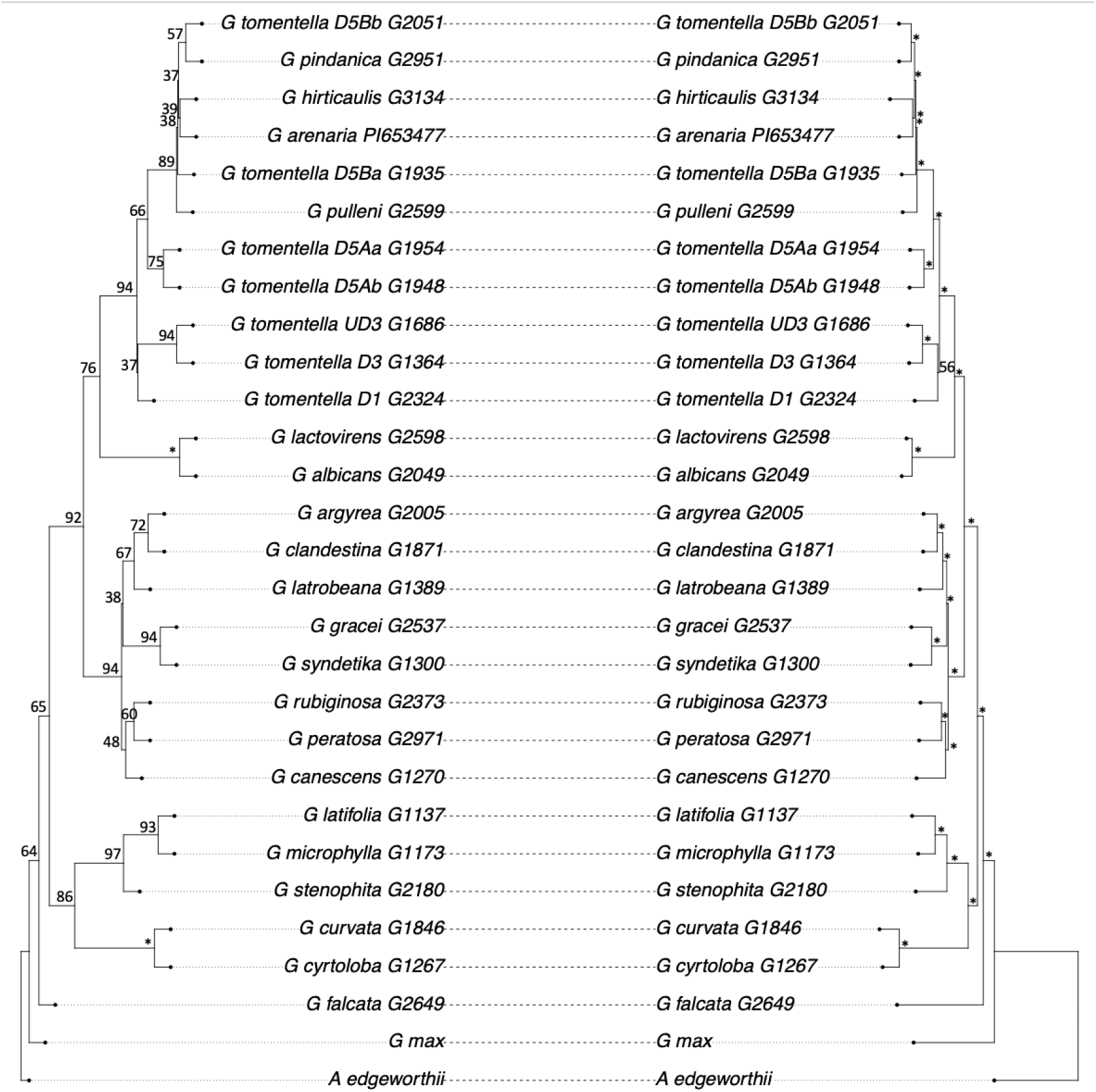
Cluster 3 cophylo ASTRAL and concatenation. Support values (quartet 1 score for ASTRAL and bootstrap for concatenation) are shown. * denotes 100% support.

**Supplemental Figure S16.**
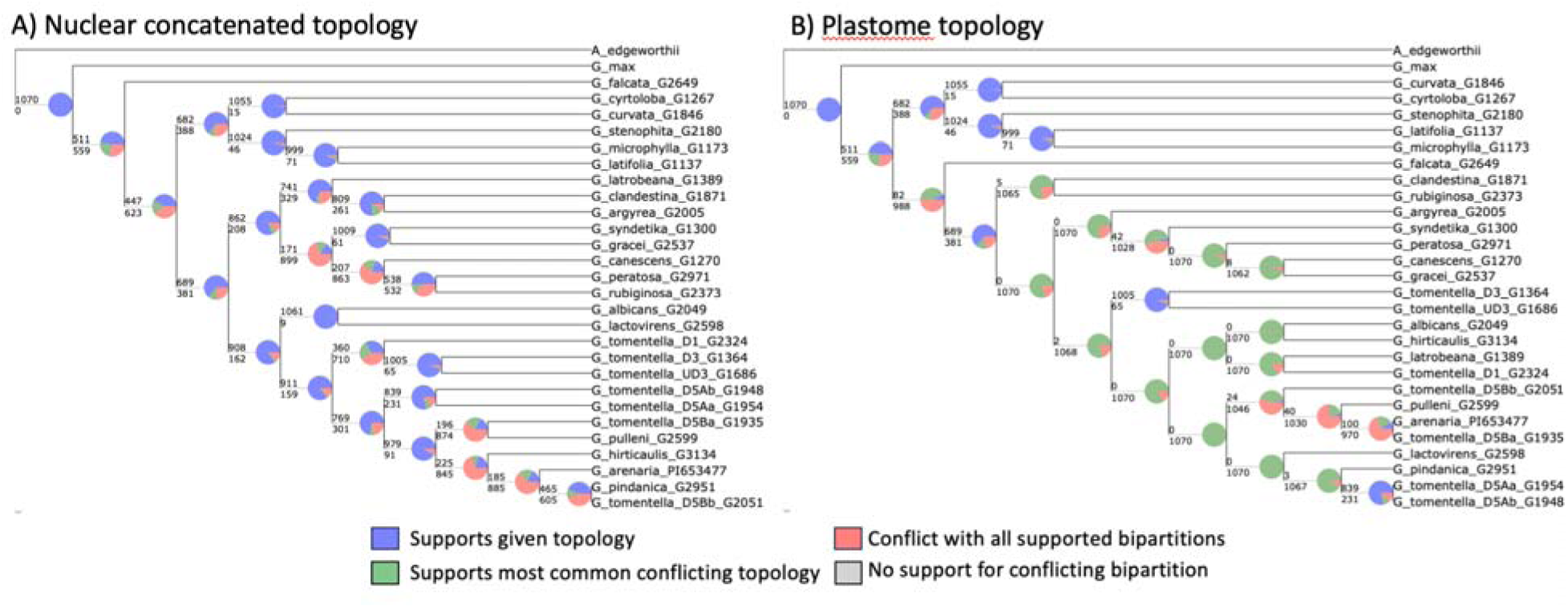
PhyParts for Cluster 1 comparing loci to the concatenated nuclear tree and the plastome tree.

**Supplemental Figure S17.**
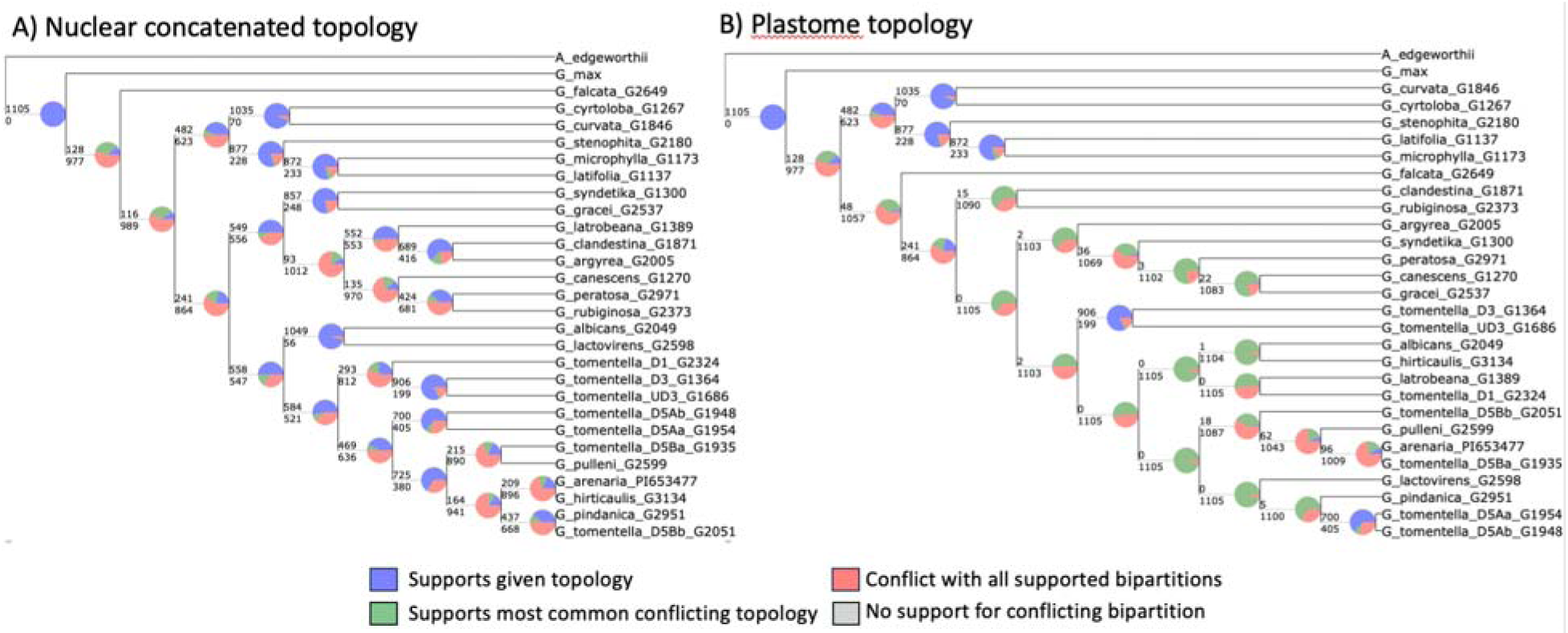
PhyParts for Cluster 2 comparing loci to the concatenated nuclear tree and the plastome tree.

**Supplemental Figure S18.**
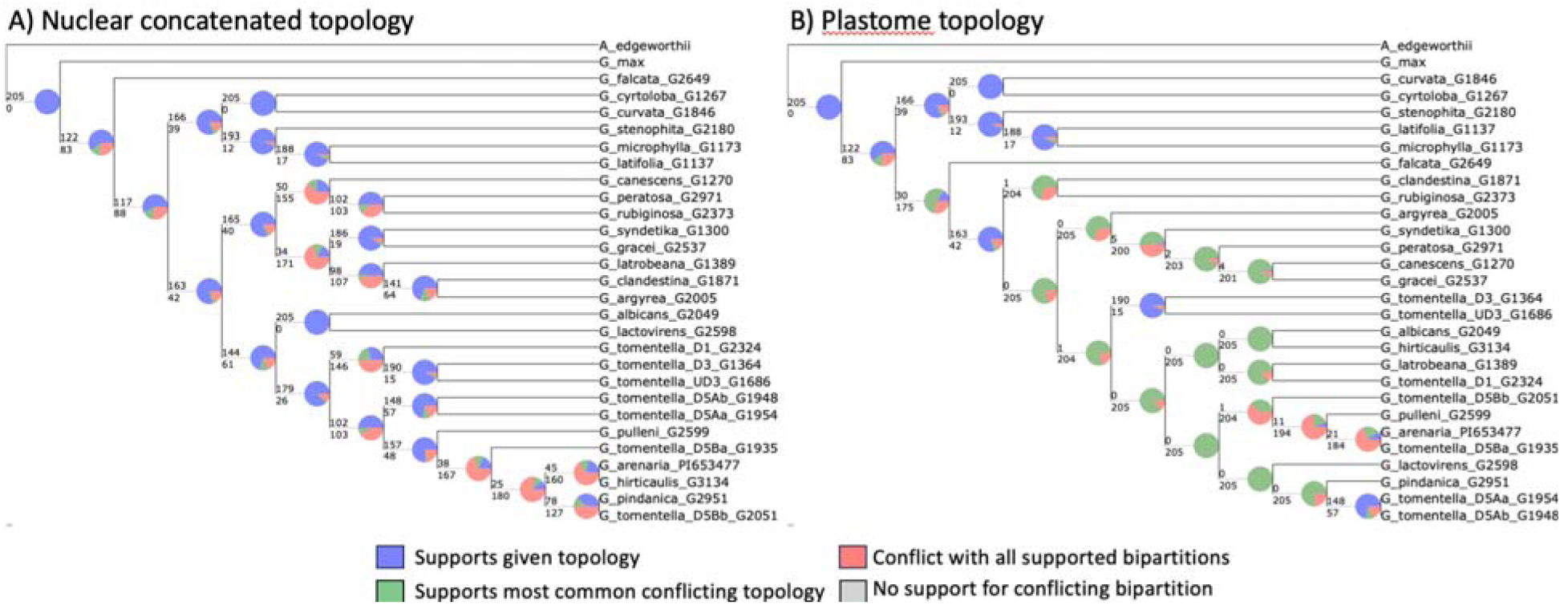
PhyParts for Cluster 3 comparing loci to the concatenated nuclear tree and the plastome tree.

**Supplemental Figure S19.**
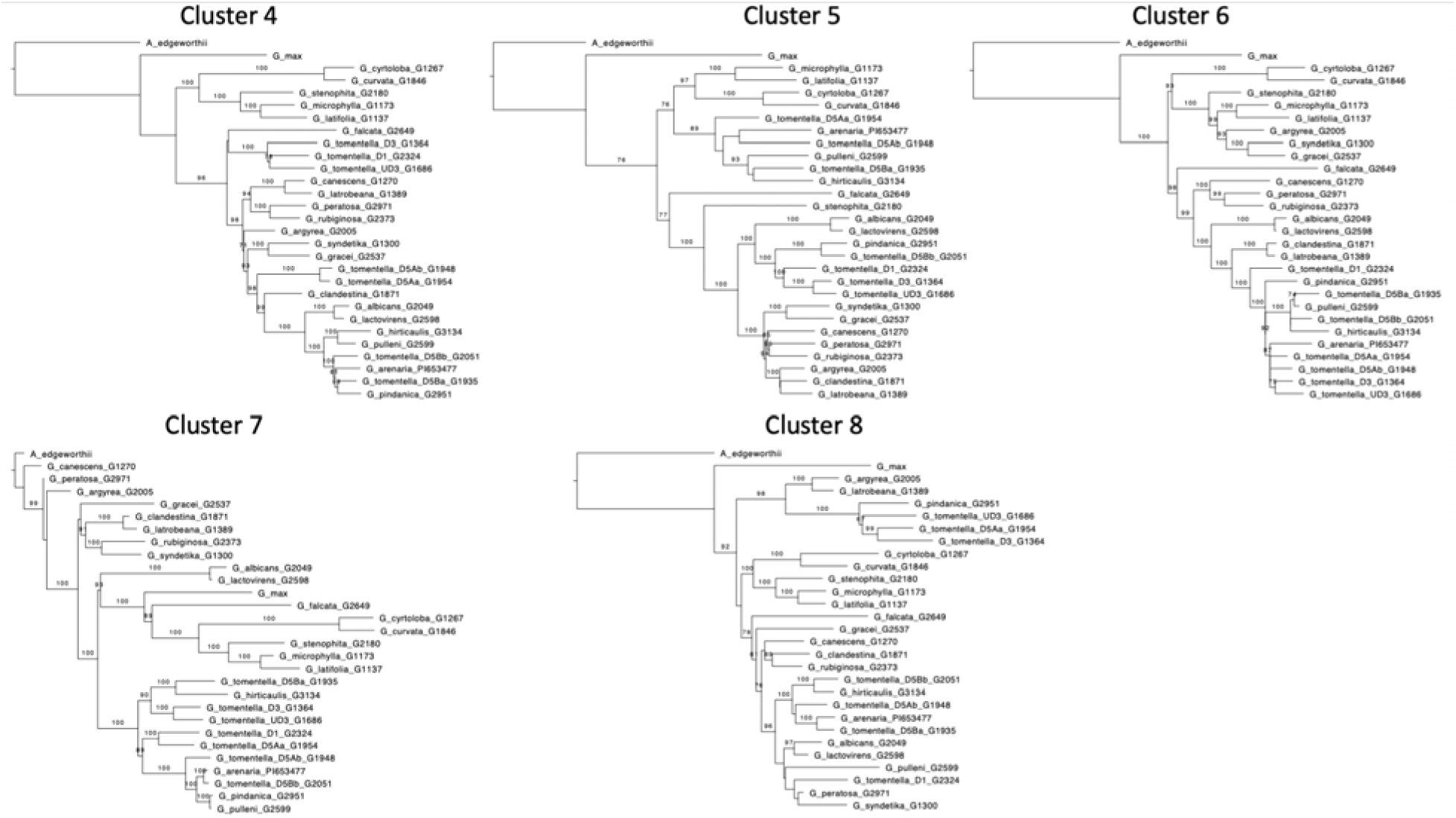
Clusters 4 to 8 represented by a unique tree topology. Bootstrap values shown for node support.

Because the two clusters identified as the best solution for this dataset did not resolve different histories as we had hoped, we explored other solutions involving more clusters. With three clusters, the same two clusters remained the dominant pattern, accounting for all but 208 trees, with 56 trees being grouped with the mostly unresolved cluster and 152 being grouped with the resolved cluster). These Cluster 3 loci included nine with evidence of recombination (one remaining after Bonferroni correction); trees were comparably resolved (avg. 18.57 nodes) and with overall similar branch lengths (avg. 0.01754, median 0.00783) to Cluster 1. Cluster 3 included the plastome tree; however, the concatenation and ASTRAL species trees reconstructed from Cluster 3 loci had topologies nearly identical to the species tree topology (Figure 1), differing only in relationships within the H-genome clade (Supplemental Figure S15-S18). Consistent with this, Cluster 3 included a higher percentage of loci with the species tree topology in the smaller Dataset 3 (18%) than did Cluster 1 (10%).

When eight clusters were recognized, these three clusters were retained and five new clusters, each comprising a single tree, were added. Trees (Supplemental Figure S19) were well-resolved (19-24 nodes), with branch lengths falling in the range of the other larger clusters. None of the loci in the new clusters showed evidence of recombination. Based on the different tests for pseudo-orthology, these five loci are not candidates for being pseudo-orthologs. The Cluster 4 tree was like the plastome tree in having *G. falcata* strongly supported as part of the A-plastome group; as in the plastome phylogeny, relationships among ADEH/HaI taxa did not follow species or genome group boundaries. The Cluster 8 tree also placed *G. falcata* within the A-plastome group but was less well-resolved. The topologies of the individual trees from the remaining clusters all had unexpected features, often moderately to strongly supported. For example, the Cluster 5 tree included a group of mostly H-genome accessions forming a clade with the B+C genome OTUs, supported at 76%; the Cluster 6 tree nested several A-genome accessions within the B-genome with 100% support; and the Clade 7 tree had to be rooted with *G. max* to make more sense, but this nested *Amphicarpaea* deeply within the A-genome, sister to *G. canescens* with 99% support.

### Utility of loci in one dataset is a weak predictor of utility in other datasets

Clearly, for Dataset 2, trees were clustered based on criteria that we do not fully understand, perhaps due to a continuum of topologies with no clear boundaries in the complexity of the enormous topology space for 27 OTUs. We therefore evaluated Dataset 2 using loci that recovered the species tree topology with the small Dataset 3. We hypothesized that one of these 141 trees should have the species tree topology as identified by both concatenation and ASTRAL for Dataset 2. We further hypothesized that the remaining 140 trees–all of them with unique topologies according to the phytools analysis–should be more like the species tree than would trees from the remaining 158 loci from the Dataset 3 analysis.

The number of resolved nodes with bootstrap support above 70% was determined for all BUSCO loci using Dataset 2. Only nine loci had all 26 nodes resolved, of which seven were among the 299 fully resolved Dataset 3 loci; however, only one of these seven loci was among the 141 Dataset 3 loci with the species tree topology. An additional 37 loci had Dataset 2 trees with 25 resolved nodes, of which 26 loci were among the 299 resolved Dataset 3 loci; 17 of these are among the 141 loci with the resolved species tree topology. The 141 Dataset 3 loci with fully resolved trees tracking the species tree had an average of 21.14 nodes resolved in Dataset 2 (compared with 15.9 nodes for all 2386 loci), with a range of 6-26 nodes resolved.

This exploration of 27-taxon topologies showed that most topologies were consistent with the topologies of the Dataset 3 analysis, notably recovering topologies similar to the species tree and plastome topologies. Disagreement among trees either involved polytomies or was confined to conflict among OTUs of clades not present in the smaller analysis. Of note was the finding that nearly all of the trees that were incorrectly rooted at the A-genome representative (*G. syndetika* in Dataset 3) were rooted at the A-genome species (*G. canescens*) in Dataset 2, with *G. canescens* being the reference used for calling SNPs. In Dataset 2 trees, the root was never placed at *G. syndetika* or any of the other A-genome species. In many (but not all) of these trees, the outgroup and *G. canescens* sequences were identical or nearly so. This problem was not encountered in loci whose Dataset 3 trees were rooted at the B-, C-, or D-genome species.

The finding that loci providing good resolution and support differed between Datasets 2 and 3, which included the same range of taxa, prompted us to analyze three 27-taxon datasets (Datasets 4-6; Table 1) composed of focused groups of taxa representing different major clades from the genus-wide sampling used in Datasets 1-3, and using all BUSCO loci. We found that the same set of loci provided very different amounts of information for different clades. Notably, although taxa in the A- and B-genomes diverged around the same time (Figure 1), far fewer loci provided resolution for relationships in the B-genome than in the A-genome (Figure 5). The number of loci with good support was more similar between the more closely related A- and DEHHaI-genome groups, but fewer than one third of the 898 loci well-resolved for either Dataset were well-resolved for both. Overall, results from the screening set (Dataset 1) were at best a weak predictor of utility for individual clades.

**Figure 5.**
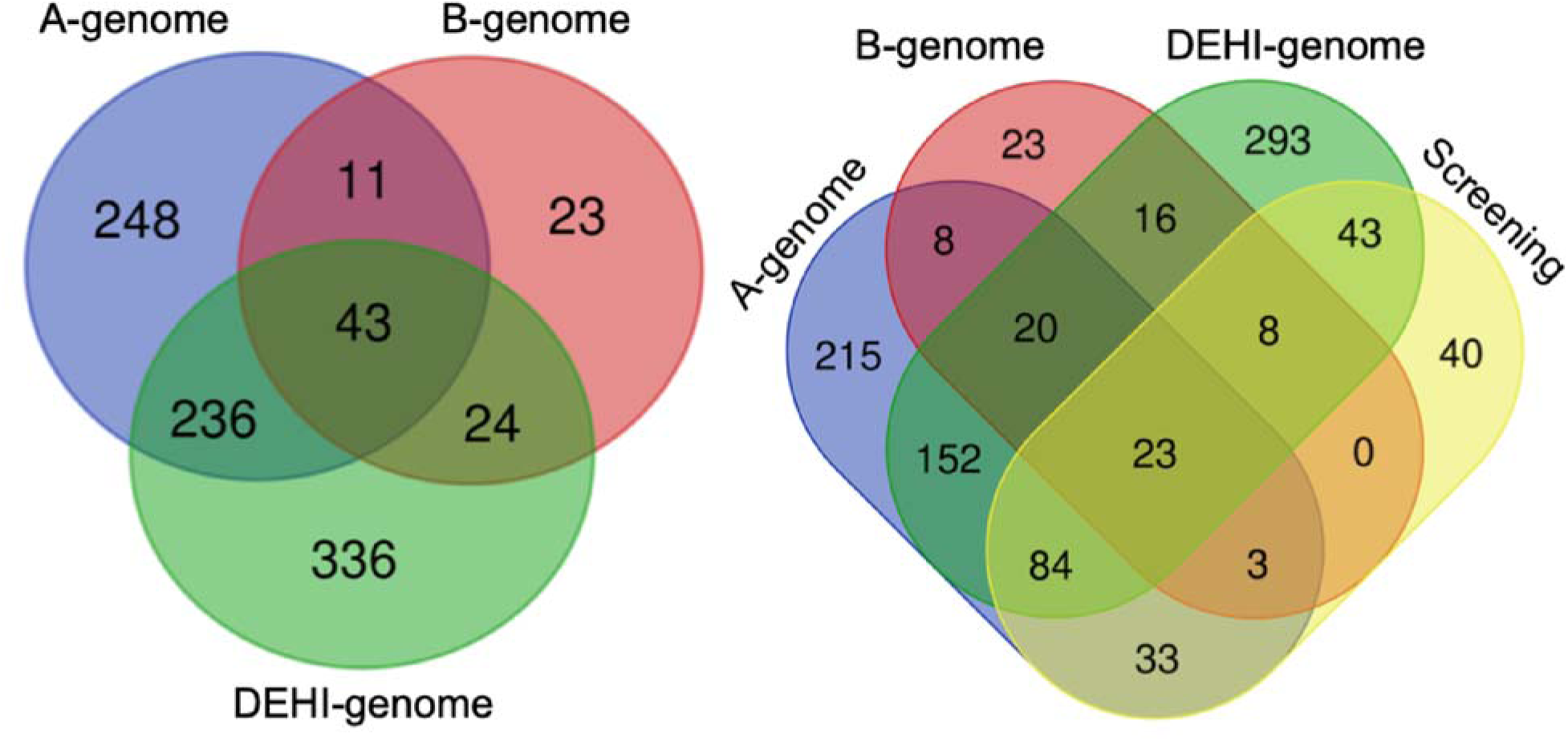
Overlap of loci that inferred phylogenetic trees with at least 23 nodes out of the 26 with bootstrap support values over 70%.

### Weak evidence for gene flow/hybridization

After a Bonferonni correction for the significance threshold (Z-score > 4.41 and p-value <1.008e-5), the ABBA-BABA analyses detected 334 cases of gene flow in the 61-taxon data set. Out of the total, 125 events involved at least two members of the same genome group (Figure 6), with 102 events involving two members of the A-genome group, 21 involving two H-genome group, and 1 each with two members of the B-genome group and Ha-genome group. Further, 46 involved a member of the A-genome group and the D-genome group with the B-genome or F-genome group, 87 events involving the B- and C-genome groups with a member of the ADEHI-genome groups. Although there were 334 significant tests for introgression in the 61-taxon data set, the D-statistic indicating the strength of gene flow was quite low, with most values ranging between 0.04 and 0.2. Pruning the sampling down to the Dataset 3 sampling, only five of the 21 ABBA-BABA events were deemed significant (Z-score > 3 with a p-value < 0.0024) with three involving the C-genome group and B-genome group with the A-, D-, and G-genome groups. The remaining two events involved the A- and D-genome groups with the F-genome group, and the C- and D-genome group with the F-genome group. The introgression events between the D-genome group and the F-genome group likely gave rise to the alternative plastome topology observed in Supplemental Figure S9.

**Figure 6.**
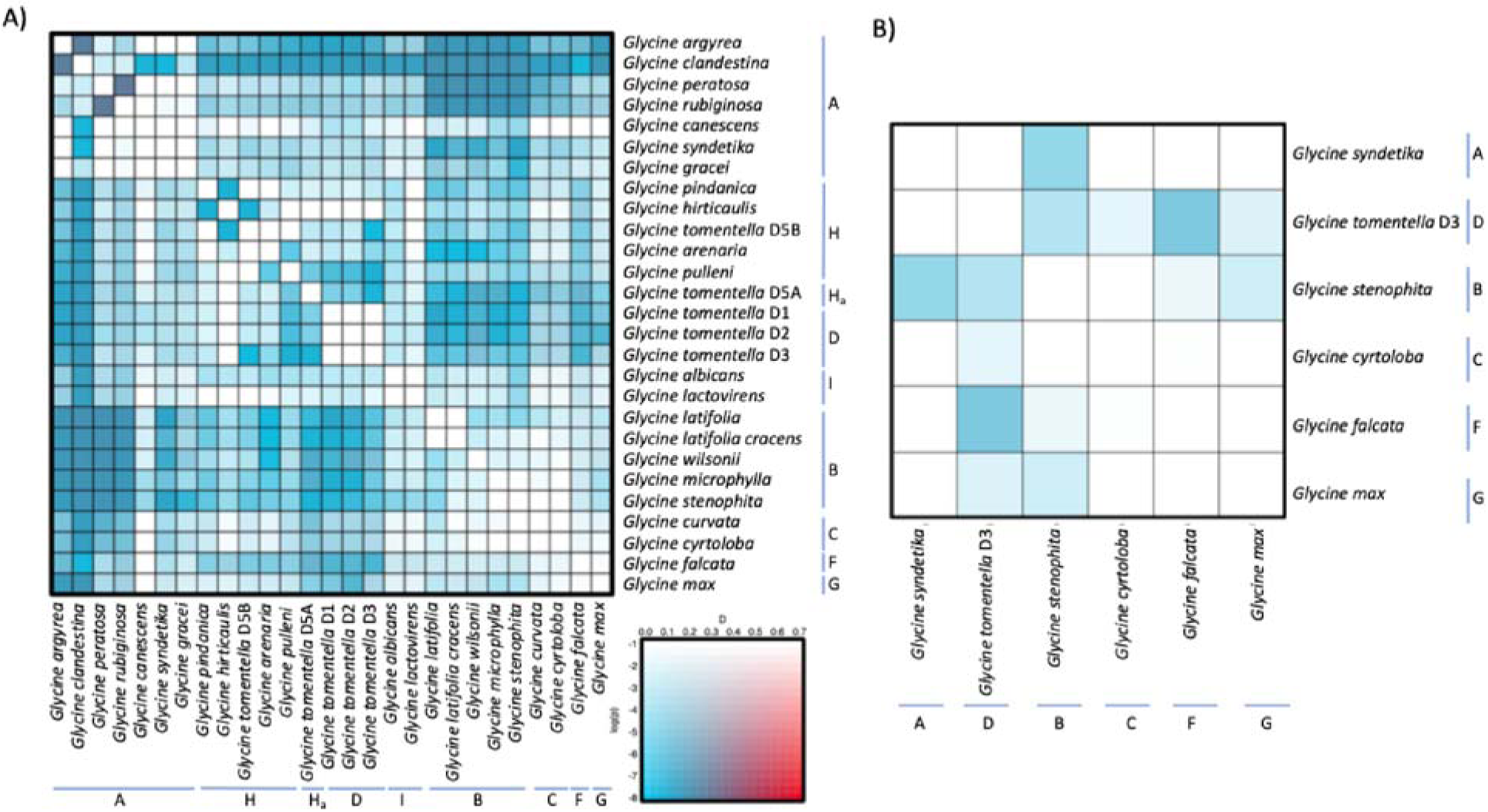
Summary of the ABBA-BABA results for the 61-taxon data set (A) and the Zhuang sampling (B) with genome groups marked. The legend denotes strength of the D-statistic and p-value for both panels.

The phylogenetic network analyses for detecting gene flow with the 299 fully resolved loci in Dataset 3 failed to find any statistical evidence for hybridization within *Glycine*. The log likelihood values for zero hybridization events was the same as when allowing up to four hybridization events.

### Pseudo-orthology is not a problem despite relatively recent Glycine polyploidy

The congruence of concatenation and ASTRAL trees suggests that ILS is unlikely to be a significant source of incongruence in Dataset 3. In contrast, pseudo-orthology could be an explanation for some or all of the nearly 50% of Dataset 3 loci with tree topologies that differed from either the species tree or the other readily interpretable topology (the plastome tree). Particularly suspect are the many incorrectly rooted trees, because in gene trees of duplicated loci that include a mixture of *Glycine* orthologues and paralogues/homoeologues, the single copy *Amphicarpaea* outgroup sequence would attach between the two paralogous clades rather than at the expected position along the branch connecting *G. max* to the perennial species.

Additionally, the correct species tree topology can be produced if the two sister clades/taxa at the base of the phylogeny are pseudo-orthologues. We therefore checked all 299 loci used in Dataset 3 for evidence of pseudo-orthology by assessing the homoeology of these loci (see Methods). Only one locus of the 141 loci with the properly rooted species tree topology was flagged as a possible case where the *G. max* gene was a different homoeologue than the other taxa (Supplemental Table S2). Because additional taxon sampling often resolves conflation of paralogy and orthology, we checked the topology of this locus in Dataset 2, and found that the tree was poorly resolved. Similar results were found with the five trees flagged as including pseudo-orthologs among the remaining 158 loci with non-species tree topologies and/or rootings. We also checked the anomalous trees identified as Clusters 4-8 (see above) and none were designated as possible pseudo-orthologs. In fact, when incorporating genomic position and collinear blocks, only 46 loci out of 2,389 (1.93%) were candidates for pseudo-orthology. When looking at the clustering in Dataset 2, 20 of the candidates fall in Cluster 2, in which there is a lack of phylogenetic signal. The remaining loci that are potentially pseudo- orthologs are more resolved, but generally in their gene trees several nodes are not well-supported. Three of the 46 loci flagged for pseudo-orthology were also noted as having significant recombination as determined with PhiPak (Supplemental Table S2)

The pseudo-orthology tests proposed by Frost et al. (2024) investigating tree length and bipartition support in gene trees from both the 1:1 orthologs and BUSCO genes showed few loci suspected of being pseudo-orthologs. For Dataset 2, 52 trees displayed a tree length larger than the mean while 1217 had bipartition support larger than the mean. The intersection of these two criteria highlighted four loci (0.17% of loci) that were candidates for pseudo-orthology (Supplemental Figure S20). Tests on Dataset 3 highlighted 16 loci with tree length larger than the average and 1265 loci showing elevated bipartition support. Only five loci (0.21% of loci) overlapped between the two metrics, none of which were highlighted in Dataset 2. Using the same metrics but for the 1:1 single copy genes identified by OrthoFinder from the genomes of Zhuang et al.(2022) we found 14 loci showing longer tree length than the average and 1366 loci showing elevated bipartition support, with 14 out of 2713 loci (0.52% of loci) overlapping between the two metrics. When incorporating the genome assemblies available in *Glycine* and collinear blocks of the BUSCO genes, none of the identified nine loci using tree length and bipartition support were likely candidates for pseudo-orthology.

**Supplemental Figure S20.**
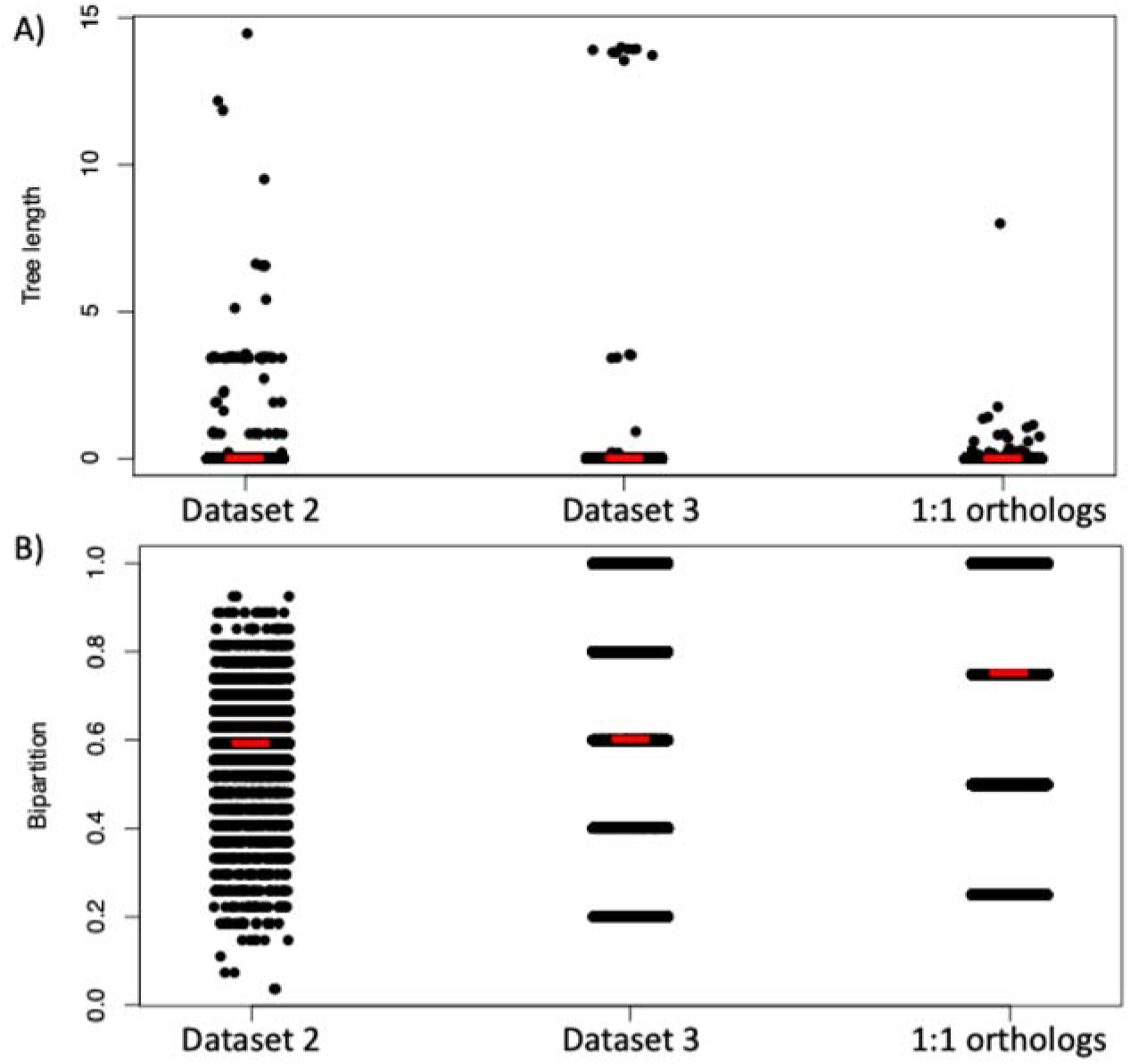
Strip plots of calculated tree length (A) and bipartition support (B) for the individual gene trees for dataset 2, dataset 3, and the 1:1 orthologs from Zhuang et al. 2022. Red lines indicate mean values, which were used as the threshold for designating problematic loci.

## Discussion

Phylogenomic analyses of over 2000 nuclear genes, the plastome, and the mitogenome clarified and extended previous phylogenetic (Sherman-Broyles et al. 2014b) and phylogenomic (Landis and Doyle 2023) studies of the ∼30 legume species comprising diploid (2*n* = 40) members of the perennial *Glycine* subgenus *Glycine*. As sister to the cultivated soybean (*G. max*) and its wild progenitor (*G. soja*), subg. *Glycine* is an important resource for improvement of a major global crop and key genetic model flowering plant. The known genome groups (Hymowitz et al. 1998) are inferred as monophyletic with strong support using both maximum likelihood bootstrap and ASTRAL normalized quartet support. However, the deepest nodes show low quartet support in the relationships between *G. falcata* and the remaining perennial species, and relationships between genome groups (Figure 1). These relationships were identified in earlier phylogenetic studies using single nuclear genes, albeit some with weak support (Sherman-Broyles et al. 2014b), and more recently in a phylogenomic study based on genotyping by sequencing (GBS) single SNP data (Landis and Doyle 2023). Two A-genome species, *G. canescens* and *G. clandestina*, known to belong to closely related species complexes (Pfeil et al. 2001) were found to be paraphyletic, and warrant further investigation.

Subgenus *Glycine* radiated rapidly in Australia around 6.5 MYA, with *G. falcata* (F-genome) sister to the remainder of the subgenus. Cytoplasmic genomes tell a similar story, but with one major exception: although the same genome groups are supported, there is a dichotomy between the (B, C) genome clade and the remainder of the subgenus, with *G. falcata* either sister to (mitogenome) or part of (plastome) the ADEHI taxa. Incongruence between the plastome and the nuclear genome has been known for some time, and was attributed to introgression rather than lineage sorting due to the shallower rather than deeper plastome coalescence (Rauscher et al. 2004). That finding is corroborated here (∼6.5 MY for F vs. remaining nuclear genomes; ∼2 MY for F vs. ADEHI plastomes). The mitogenome of *G. falcata* is strongly supported as joining an A-mitogenome clade, but the coalescence date is considerably older (∼5 MY), although less than the divergence with the (B, C) mitogenome clade (∼6 MY). The mitogenome tree is overall less well supported than the plastome tree, but dating and topological differences between the two warrant further investigation, as incongruence between mitogenome and plastome has been observed across land plants (Tyszka et al. 2023). In any case, the cytoplasmic history for subgenus *Glycine* differs from that of the species tree. Consequently, we searched for a signal of this history in nuclear gene tree topology space.

We found that the nuclear gene tree topology space for *Glycine* is flat when more than a handful of taxa are analyzed; this space is a sparse, featureless landscape in which every gene has a unique topology. In all but the smallest dataset there is no peak of loci with the species tree topology, let alone secondary peaks representing competing phylogenetic signals. The distribution of these singular trees is also unstructured, and no biologically meaningful patterns could be discerned by clustering. Yet these incongruent trees, even in relatively small subsets, contain sufficient information to permit the reconstruction of a robust species tree and to find the signal of the cytoplasmic introgression event in the nuclear genome. Notably, even the ∼1000 loci that cluster together based on their low phylogenetic informativeness (Cluster 2 in Dataset 2) contain sufficient information for reconstructing an accurate species tree using ASTRAL, and an even more robust tree using concatenation.

That *Glycine* species comprise “collections of genomic regions with different histories” (Rosenberg and Nordborg 2002) is not a surprise, of course. Rokas et al. (2003) inaugurated a new phylogenomic era when they sampled 106 nuclear loci from the complete genomes of seven *Saccharomyces* yeast taxa and a *Candida* outgroup; these had over 20 different topologies but produced a single robustly supported supergene tree when concatenated. In a *Nature* editorial on this breakthrough paper provocatively titled “Ending Incongruence”, Gee (2003) concluded that although Rokas et al. (2003) had “raised the game of phylogenetic reconstruction to a new level”, the true species phylogeny is unlikely ever to be known, and this should be a “source of unease” for evolutionary biologists. Indeed, using the same genes with the larger yeast dataset of Rokas and Carroll (2005), Jeffroy et al. (2006) reported that each of the 106 gene trees had a different topology; they concluded that phylogenomics had in fact ushered in the “beginning of incongruence”. Such studies corroborated longstanding population genetic theory positing that neutral evolutionary phenomena (specifically deep coalescence/ILS) are expected to cause unlinked loci to have incongruent gene trees (e.g., Nei 1987), which has become a fundamental assumption of phylogenomics (Edwards 2009; Bravo et al. 2019).

What seems more unexpected is how fragmented histories are in *Glycine*–there is no major species tree signal in gene tree topology space for even the relatively small (27 or 61 taxon) datasets we analyzed. Again, this is not unprecedented, as Jeffroy et al. (2006) reported for the 106 genes in 14 yeast species, but this phenomenon seems mostly ignored in the literature. Presumably this is because, despite the exhortation of Bravo et al. (2019) to “embrace the many heterogeneous genomic signals” emerging from phylogenomic analyses, the focus of most researchers is to identify “the” species tree. When this is the goal, gene trees can be considered a nuisance parameter (e.g., Heled and Drummond 2010). Perhaps for that reason, although there has been interest in exploring the differential utility of genetic loci for phylogenetic analysis using both concatenation and coalescent approaches (e.g., Mirarab et al. 2014; Mongiardino Koch 2021; Lozano-Fernandez 2022), there has been much less interest in the topologies of the gene trees themselves. Although Bravo et al. (2019) state that deciphering the diverse histories that produce “many heterogeneous genomic signals” is “one of the grand challenges of evolutionary biology”, they do not discuss how incongruence is structured in gene tree topology space.

Diverse histories do not occur as collections of loci with shared topologies standing out against an otherwise flat landscape, dominated by the species tree topology. Why is this? Consider a dataset of 100 unlinked, internally non-recombining loci (c-genes) sampled from three taxa and an ingroup. If all loci produce fully resolved gene trees, neutral expectations are that the species tree will be the most common topology; Maddison (1997) showed an example in which 75% of gene trees track the species phylogeny, with the remaining trees equally split between the two ILS alternatives. A second speciation event would add an OTU, increasing tree space, with the same expectation of the species topology being the most common of the three possible topologies. But assuming the same demographic and speciation parameters, again only 75% of the trees will correctly track the species tree, meaning that now the species tree topology is represented by only 57 loci, with the remaining 18 trees split between two new topologies, each having the species tree topology at the basal node but ILS alternatives at the tips. Meanwhile, the ILS peaks from the first speciation event experience the same reduction in number of supporting loci; the non-species tree first node of each subtends a more distal node consistent with the species-tree, with dispersion of two new topologies into gene tree topology space, each having two successive nodes tracking ILS histories. This flattening process is accelerated if there are other historical signals in addition to ILS; for example, introgression will also “steal” loci from other peaks (notably the species tree), dispersing loci to previously unoccupied locations and further flattening topology space. The number of gene trees can never exceed the number of loci, so as additional speciation events occur the 100 gene trees are dispersed in an exponentially expanding tree space. Eventually each locus will have a unique topology. When gene tree topology space is flat, at most one locus can have the complete species tree topology, and even this is highly unlikely.

Thus, for a given set of taxa, individual loci have singular histories; but rather than one locus having a species tree history, another an ILS history, and a third an introgression history, the tree of each locus comprises fragments of many different histories shared in unique combinations with other loci. These histories necessarily shift with the set of taxa included in the analysis; loci themselves have properties (Shen et al. 2016b; e.g., evolutionary rate, base composition) that may make them generally more, or less, useful for constructing a gene tree, but for the most part utility is taxon-dependent, as we found for 27-taxon datasets sampling different clades of closely related *Glycine* species (Datasets 2, 4-6) and even when comparing datasets composed of different numbers of the same set of taxa (Dataset 2 vs. Dataset 3).

Fortunately, phylogenomic methods do an excellent job of extracting historical signals dispersed among different loci and assembling them into a species tree. Concatenation does so by ignoring gene trees entirely and relying on individual characters to track history, whereas pseudo-coalescent methods such as ASTRAL succeed by considering gene trees one node at a time and relying on the fact that the species tree is always the most common topology (Pamilo and Nei 1988; Kubatko 2019; Mirarab 2019). However, if the strongest signal–that of the species tree–can only be detected in the flat landscape of gene tree topology space by aggregating fragments of the overall history from many loci, it stands to reason that competing histories will be more difficult to detect. In *Glycine*, we had a hypothesis to test because of the striking incongruence of the plastome and mitogenome with the species tree, detectable thanks to the unique ability of these large, character-rich single coalescent genes (Doyle 2022) to provide detailed phylogenies. This allowed us to use PhyParts on the plastome topology to identify the relatively small number of nuclear loci that track, at the key node, the putative introgression. Without such a hypothesis, identifying heterogeneous signals in the data would be little more than a fishing expedition. If the vision of Bravo et al. (2019) is to be realized, better methods need to be developed to identify biologically interesting non-species tree signals fragmented and scattered across the nuclear genome.

Pseudo-orthology, produced by conflation of paralogous and orthologous loci, has been cited as a potential problem since the dawn of molecular phylogenetics (Fitch 1970; Doyle 1992; Maddison 1997; Edwards 2009). It could be a particular problem for *Glycine* species, whose genomes were shaped by a recent (>13 MY) ancestral polyploidy event, from which one copy has been lost from over 25% of homoeologous pairs (Schmutz et al. 2010). This means that thousands of loci are candidates for the kind of alternate loss of duplicate copies required for assembling phylogenetic datasets consisting of a mixture of orthologous and paralogous genes (Smith and Hahn 2022). Although modeling studies have shown that the probability of sampling pseudo-orthologs is low relative to the probability of orthologs, and that the impact of inclusion of pseudo-orthologues on multi-gene analyses should be minimal (Smith and Hahn 2022), pseudo-orthology continues to be a source of concern in the phylogenomics era (e.g., “artifactual orthologs” of Frost et al. (2024). Our results, based on mapping 2386 loci to sequenced genomes representing the major *Glycine* lineages (Zhuang et al. 2022) show that the fraction of loci affected by pseudo-orthology is vanishingly low. These reassuring findings agree with the theoretical conclusions of Smith and Hahn (2022). The modular, fragmented nature of phylogenetic signal also means that even in a tree distorted by pseudo-orthology most nodes will contribute to the correct species tree. This is yet another reason why gene families are useful for phylogeny reconstruction (Smith et al. 2022).

Our attempts to identify clusters of species tree-like and plastome tree-like topologies in gene tree topology space failed, despite using methods designed to identify related topologies (Smith 2022), including Robinson-Foulds distances that have been found to be useful for clustering gene trees (Kuhner and Yamato 2015; Smith 2020; Spirin et al. 2024). Perhaps this is a result of the overall similarities of the topologies among these closely related species. The complex pattern of relationships produced by lineage sorting and other sources of incongruence could create a network of similarities too interdigitated to permit discrimination of discrete regions of this small portion of tree space. How the flattening and dispersion of locus topologies during speciation populates gene tree topology space would be interesting to investigate further.

Our results make very clear the enormous improvement of phylogenomics over phylogenetics for identifying organismal histories. There is no longer a need to choose between investing in taxon vs. gene sampling (Rokas and Carroll 2005) both of which mitigate many problems, including pseudo-orthology (Smith and Hahn 2022). But even more significantly, the unrecognized flatness of gene tree topology space made it inevitable that when only a few loci could be sampled, their gene trees would be incongruent, leading to ad hoc explanations and uncertainty about the species tree. The ability to extract diverse signals from numerous loci and reassemble those signals into a single tree summarizing taxon relationships is a tremendous advantage, providing a hypothesis against which other biologically meaningful signals can be tested.

## Data availability

Raw reads underlying BUSCO genes were deposited in the SRA with accession BioProject (Provided upon acceptance), see Supplemental Table 3 for accessions numbers representing specific accessions. Alignments for each locus with the different taxonomic sampling have been submitted to Dryad.

## Acknowledgments

This work was supported by the US Department of Agriculture National Institute of Food and Agriculture Hatch projects 1014310 and 7002754 to J.J.D. We thank K. Donohue, L. Shi, T. Kelliher, and G. Wu for laboratory and personnel support, and and L. Mumm, J. Ho, S. Miles, K. Jones, K. Williamson, and S. Kanjilal for licensing and guidance on intellectual property.

## Supplemental Tables

Supplemental Table S1. Functional description, BUSCO ID, and associated *G. max* sequence for BUSCO loci used for phylogenetic inference.

Supplemental Table S2. Summary of each locus used, including number of sites, number of conserved and variable sites in the 61-taxon alignment, informative sites, and missing data. Also included are which cluster the locus part of in Dataset 2 clustering and whether the locus was a candidate for pseudo-orthology.

Supplemental Table S3. Summary of the 61 accessions used for phylogenomic analyses and associated SRA numbers.

Supplemental Figure S1. Maximum likelihood tree of 570 *Glcyine* accessions using 100 genes; the first five BUSCO genes on each of 20 chromosomes. Blue boxes are loci used for the 61 accessions making up Dataset 1.

---

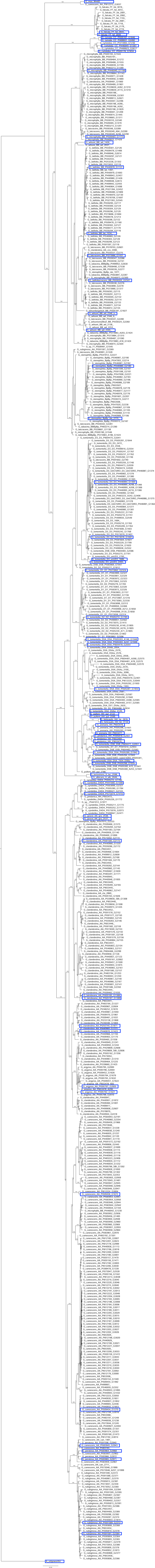

## Notes

### Competing Interest Statement

The authors have declared no competing interest.

